# A general theoretical framework for trait-based eco-evolutionary dynamics: population structure, intraspecific variation, and community assembly

**DOI:** 10.1101/2021.10.01.462789

**Authors:** Jonas Wickman, Thomas Koffel, Christopher A. Klausmeier

## Abstract

To understand how functional traits shape ecological communities it is necessary to understand both how traits across the community affect its functioning and how eco-evolutionary dynamics within the community change the traits over time. Of particular interest are so-called evolutionarily stable communities (ESCs), since these are the end points of eco-evolutionary dynamics and can persist over long time scales. One theoretical framework that has successfully been used for assembling ESCs is adaptive dynamics. However, this framework cannot account for intraspecific variation— neither locally nor across structured populations. On the other hand, in moment-based approaches, intraspecific variation is accommodated, but community assembly has been neglected. This is unfortunate as some questions regarding for example local adaptation vis-a-vis diversification into multiple species requires both facets. In this paper we develop a general theoretical framework that bridges the gap between these two approaches. We showcase how ESCs can be assembled using the framework, and illustrate various aspects of the framework using two simple models of resource competition. We believe this unifying framework could be of great use to address questions regarding the role of functional traits in communities where population structure, intraspecific variation, and eco-evolutionary dynamics are all important.

## Introduction

In recent years, the need to understand intraspecific variation of functional traits has been increasingly recognized as important for understanding the functioning of ecological communities (Albert et al., 2010; Violle et al., 2012). For example, intraspecific variation has been shown to be important in relation to niche differentiation and environmental filtering (Paine et al., 2011) and pairwise species interactions (Kraft et al., 2014). Such intraspecific variation in functional traits has also been shown to often be substantial compared to variation between species (Siefert et al., 2015).

If species have heritable variation in their traits, natural selection will act on that variation. In order to understand these effects, theorists and modelers have used models in quantitative genetics (Lande, 1979; Lande and Arnold, 1983) and moment-equation-based frameworks (Wirtz and Eckhardt, 1996; Norberg et al., 2001; Savage et al., 2007) where this variation and selection is taken into account. These types of models have been used to evaluate how intraspecific variation influences species coexistence (Barabás and D’Andrea, 2016) and how spatial gradients interact with standing variation (Norberg et al., 2012). However, these frameworks typically do not account for mutations, diversification into multiple species is hard to accommodate, and robust methods for determining whether communities are evolutionarily stable with regards to the addition of further species do not exist. On the other hand, another prominent framework for modeling eco-evolutionary dynamics known as adaptive dynamics (Metz et al., 1992; Dieckmann and Law, 1996; Geritz et al., 1998; Dercole and Rinaldi, 2008) readily accommodates questions of diversification and evolutionary stability. However, it relies on the assumption that species have no intraspecific variation in their traits, so it cannot be used to understand the eco-evolutionary consequences of intraspecific trait variation.

Over the last decade, starting with Sasaki and Dieckmann (2011), a new framework has emerged to try to bridge this gap and study problems that require including facets of both moment-based and adaptive-dynamics-based eco-evolutionary frameworks. Although several aspects of the theory such as evolutionary branching (Sasaki and Dieckmann, 2011), and the effects of multivariate traits (Débarre et al., 2014) have been studied, much still remains to be worked out. In particular, general theoretical and modeling tools for assembling evolutionarily stable communities while still incorporating intraspecific variation are not yet available. Such tools would be valuable, as evolutionarily stable communities can serve as good model communities as they can persist over long time scales (Edwards et al., 2018), and incorporating intraspecific diversity would allow us to explore how different ecological conditions affect the total diversity of traits in assembled communities. Furthermore, these eco-evolutionary models taking both multiple species and intraspecific variation into account have so far almost exclusively neglected population structure (but see Débarre et al., 2013, for an example with spatial structure). The combination of multiple species and structured populations is interesting as modeling the joint trait distribution across species and spatial patches could help address questions of how diversity is structured under the influence of evolution in complex communities where both local adaptation within species as well as separate species occupying different parts of trait space are possible.

In this paper we synthesize many recent advances in describing eco-evolutionary dynamics through moment equations taking the total density, mean trait vectors, and variance-covariance matrices into account, while also accounting for phenotypic mutations from first principle. Furthermore, we add some key missing components for using this theory for eco-evolutionary modeling, in particular techniques for assembling eco-evolutionarily stable communities. We derive generic moment equations for class-structured populations, which can then easily be adapted to specific examples, and we showcase two such example to illustrate various facets of the modeling framework.

## Methods and models

In this section we will specify our generic model for class-structured communities, show how moment approximations can be derived, and devise a community-assembly procedure to assemble eco-evolutionarily stable communities. We will illustrate the generic models with a simple two-patch model of resource competition.

### Generic class-structured model and moment equations for fixed number of species

We consider a community of clonally reproducing individuals structured in *K* classes (e.g., spatial patches or age classes), where the rates of the ecological processes for individuals in the community are governed by an *n*-dimensional trait vector *ξ* = (*ξ*_1_, *ξ*_2_, …, *ξ_n_*) ∈ ℝ^*n*^. Each component of the trait vector encodes some quantitative trait of the organism, e.g., body mass or degree of resource specialization, with *n* different traits being tracked in total. The trait-density distribution in each class *k* is given by *v_k_*(*ξ*), *k* = 1,2,…,*K*, which describes how abundant individuals with a given trait vector in class *k* are. Figure 1 shows examples of trait-density distributions in one- and two-dimensional trait space. We let **v**: = (*v*_1_,*v*_2_, …,*v_K_*) be the vector of density distributions over all the classes.

**Figure 1:**
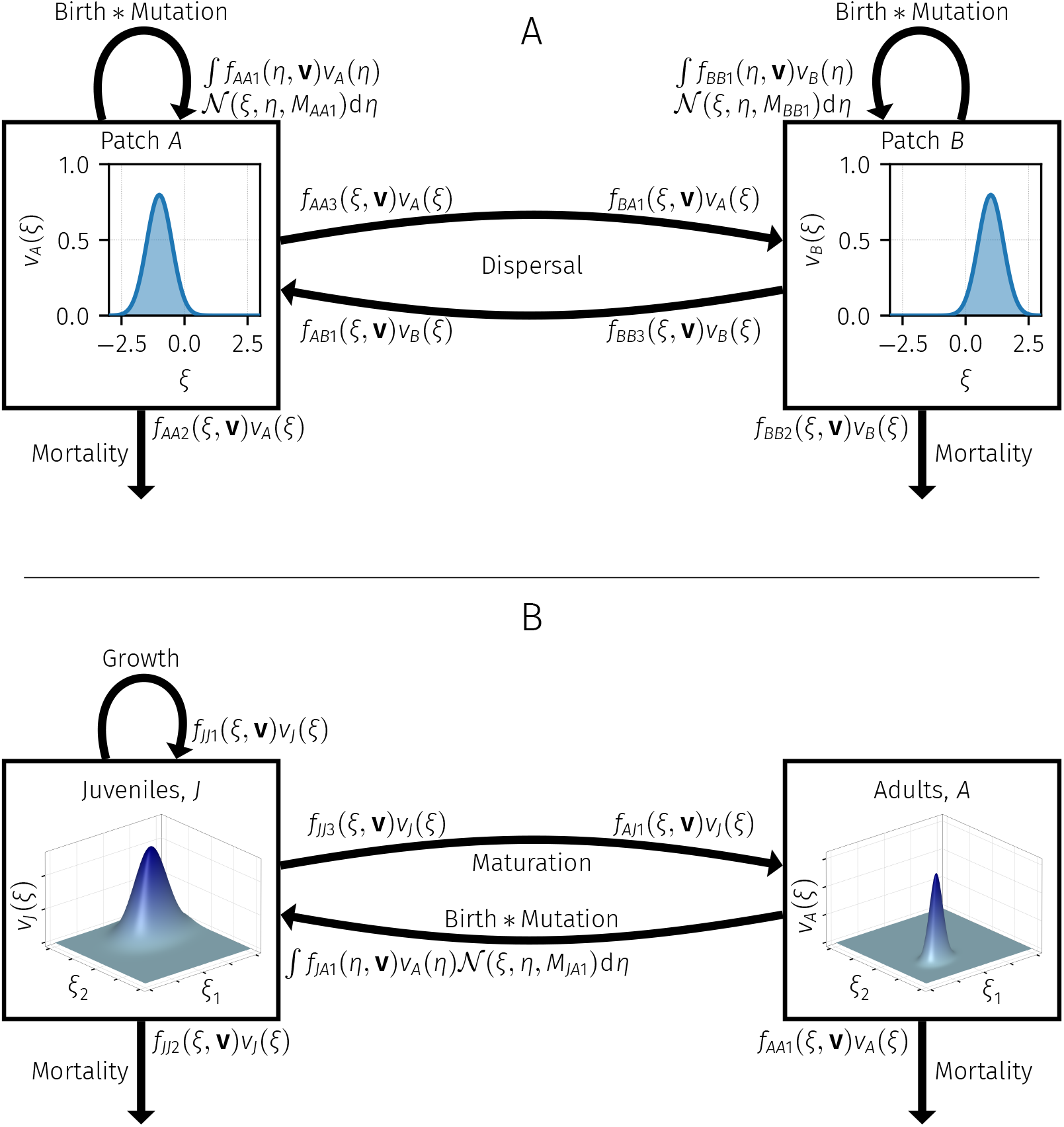
Two illustrations of the generic class-structured trait-space equations. Rate functions placed at bases of arrows indicate losses and rate functions placed at heads of arrows indicate gains. All per capita rate functions *f_klm_* can depend both on the trait of the individual under consideration, *ξ*, as well as the trait-density distributions in all classes **v**. **(A)** A two-patch model with local birth and mortality on each patch and dispersal between the patches with one trait. New individuals are born at (per capita) rates *f*_*AA*1_ and *f*_*BB*1_ on each patch respectively, and the traits of the newborn individuals are filtered through a convolution with normal distributions 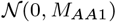 and 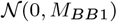 with zero means and variance-covariance *M*_*AA*1_ and *M*_*AA*2_ due to mutations on each patch respectively. Local mortality on each patch occurs at rates *f*_*AA*2_ and *f*_*BB*2_ respectively. Individuals disperse out of patch *A* at rate *f*_*AA*3_ and arrive at patch *B* with rate *f*_*BA*1_. Conversely, individuals disperese out of patch *B* at rate *f*_*BB*3_ and arrive at patch *A* at rate *f*_*AB*1_. **(B)** A two-stage model with juveniles and adults with two traits *ξ* = (*ξ*_1_, *ξ*_2_). Juveniles grow at rate *f*_*JJ*1_ and suffer mortality at rate *f*_*JJ*2_. Juveniles mature out of the juvenile stage at rate *f*_*JJ*3_ and into adults at rate *f*_*AJ*1_. Adults suffer a mortality at rate *f*_*AA*1_ and produce new juveniles at rate *f*_*JA*1_. The new juveniles thus produced have thier traits filtered through a convolution with a normal distribution 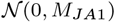 with mean zero and variance-covariance matrix *M*_*JA*1_ due to mutations.

The trait-density *v_k_*(*ξ*) in each class changes over time in response to various ecological processes. For each focal class *k* and source class *l* we specify *N_kl_* ecological processes with per-capita rates *f_kim_*(*ξ*, **v**) indexed by *m* = 1, 2, …,*N_kl_*. For within-class processes (*l* = *k*) these processes may include for example local birth and mortality rates on a spatial patch. For between-class processes (*l* ≠ *k*) they may describe for example how *v_k_* changes in response to immigration from spatial patch *l*, or from births from adult class *l* if *k* is a juvenile class in a stage-structured model. We also consider the possibility that each such process can have replicative infidelity associated with it, so that a phenotype with trait vector *η* undergoing the process with rate *f_klm_* will yield a normal distribution of individuals. Write 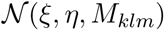 for the probablity-density function with argument *ξ* of a normal distribution with mean vector *η* and variance-covariance matrix *M_kim_*. For simplicity, we shall refer to *M_kim_* as the mutation variance-covariance matrix. The instantaneous rate of change to individuals with trait due to the ecological process with rate *f_klm_* and mutation variance-covariance matrix *M_klm_* is then given by the convolution 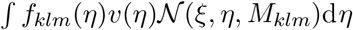. For processes with no mutation—which will typically be the case, except for births—we set *M_klm_* = 0 and interpret it in the delta-dirac sense so that

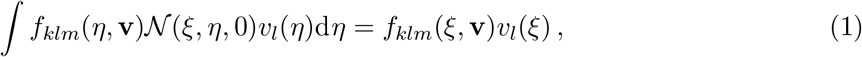

Summing over all classes and processes yields the complete equations

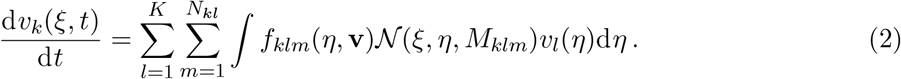

We shall refer to these types of equations as *trait-space equations.* As in adaptive dynamics and trait-diffusion approaches (Merico et al., 2014; Le gland et al., 2020; Nordbotten et al., 2020), we are not here concerned with any specific genetic makeup that would result in these dynamics for phenotypic traits, but simply assume that new heritable variation is generated in some ecological process such as births and mutations. For a single class, our setup is however very similar to the one-locus continuum-of-alleles class of models (Kimura, 1965; Bürger, 1986). Our equations also have various things in common with other eco-evolutionary approaches; we discuss these similarities further in the discussion section.

In Fig. 1 we illustrate two example models in the framework of Eq. 2: a two-patch model with a single scalar trait and a stage-structured model with one juvenile and one adult stage with two traits. We will work out the two-patch example in Box 1, and the stage-structured model is treated in section “Additional example: A stage-structured resource-competition model”. In section “Traitspace equations” of Box 1 we show an example of trait-space equations for the two-patch model, and how it relates to the generic trait-space equations (Eq. 2).

In general, Eq. 2 will not be analytically tractable; even the equilibrium of Eq. 2 will in general be hard to ascertain, even for very simple examples with only a single class (e.g., Kimura, 1965; Bürger, 1986). Numerically too, discretizing trait space and solving Eq. 2 requires significant computational power; for multiple traits especially, the feasibility of numerical explorations of the equations is highly limited. Thus, rather than solving the trait-space equations directly, we instead derive moment equations that tracks the total density, mean trait, and variance-covariance of several peaks approximating the trait-density distribution in each class. Our derivations closely follow similar methods in earlier work (Wirtz and Eckhardt, 1996; Norberg et al., 2001; Bruggeman, 2009; Merico et al., 2014; Sasaki and Dieckmann, 2011; Débarre et al., 2014; Nordbotten et al., 2020), but differs in considering class-structured populations, adding new variation through a mutational kernel, and multiple traits simultaneously.

We assume that the densities *v_k_*(*ξ*) can be usefully split into *S* peaks (which we will refer to as species) so that 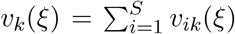, and that each of these peaks can be approximated by a normal distribution so that

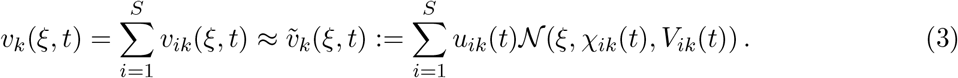

Here *u_ik_* = ∫ *v_ik_*(*ξ*)d*ξ* is the total density of species *i* in class *k*, 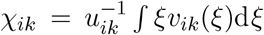 is the mean-trait vector of species *i* in class *k*, and 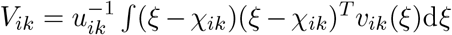 is the variancecovariance matrix of species *i* in class *k*. We also let 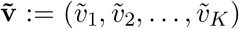 be the vector of all approximate trait distributions over all classes. Furthermore, we make the approximating assumption that each species is reproductively isolated, so that all transitions can be computed species-by-species without any trait-density ‘leaking’ between species. For now, we take the number of species *S* to be arbitrary; we will address the question of how to determine *S* in section “Eco-evolutionary community assembly”.

We define the per capita population rate of process *f_klm_* for a population with mean trait *ξ* and variance-covariance *W* as

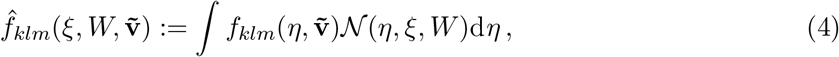

which averages the trait-dependent per capita rates over the normal phenotype distribution. Then, under these assumptions the moment equations approximating Eqs. 2 for species *i* = 1,2, …,*S* are then given by (see Appendix A for the derivation)

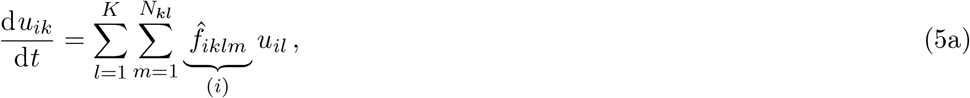

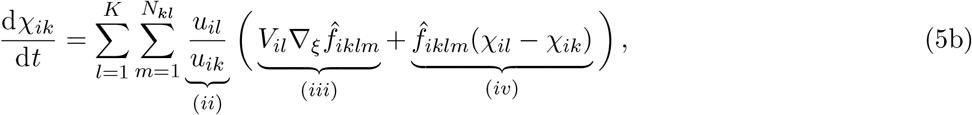

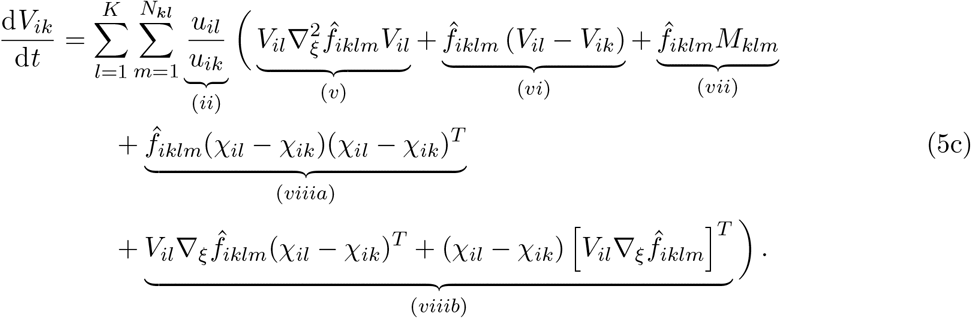

Here, the superscript *T* denotes vector or matrix transpose. Furthermore, the arguments of all rate functions have been suppressed for notational clarity, and

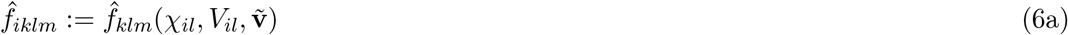

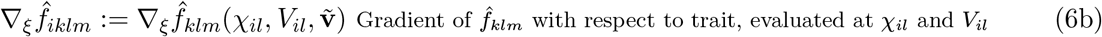

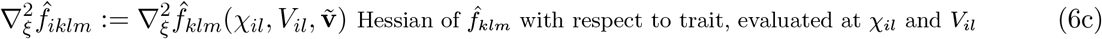

The integral in Eq. 4 is analytically tractable for some functions—for example, exponentials, gaussians, and polynomials in the scalar-trait case (see e.g., Owen, 1980)—but is in general not solvable directly. In such cases, assuming that the variance-covariances of the species are not too large, we can Taylor-expand the process rate functions *f_klm_* around the mean traits to some order. In particular, to third order this yields

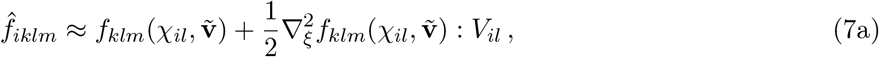

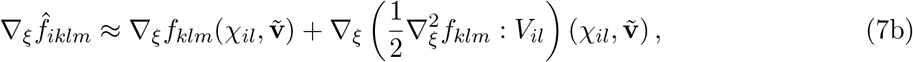

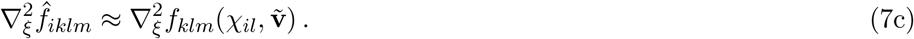

where “:” is the double dot product, which for two square *n × n* matrices *A* and *B* is given by 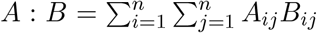. We shall use this third-order approximation for the remainder of this text. See Appendix A for expressions of arbitrary order for the Taylor expansions.

As noted in other moment-based frameworks (e.g., Norberg et al., 2001), in addition to being more tractable, moment equations can also be more interpretable than the trait-space equations from which they are derived. We now go through the various terms and factors in these equations and their interpretations. Figure 2 illustrates how these terms come about through the dynamics of the trait-space equations and their effects.

**Figure 2:**
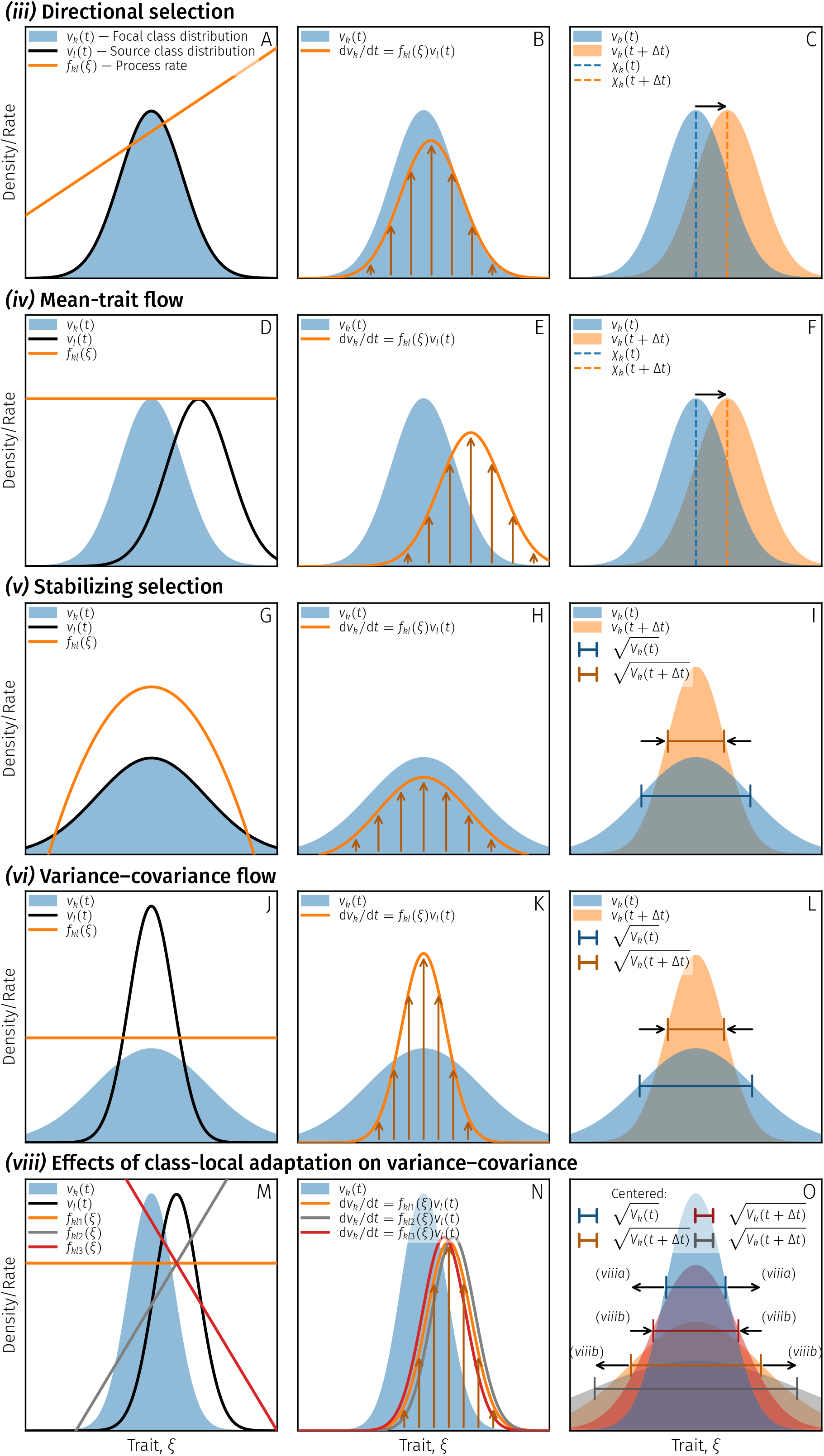
**Illustration of terms** (*iii*), (*iv*), (*v*), (*vi*), **and** (*viii*) **of Eqs. 5**. For visual clarity and purposes of explanation, these panels are mock-ups and are not true calculations based on the trait-space equations. **(A–C)** Causes and effects of term (*iii*) of Eqs. 5. Panel A shows trait-density distributions *v_k_*(*ξ,t*) and *v_l_*(*ξ,t*) of focal class *k* and source class *l* at time *t* along with a rate function *f_kl_* describing how *v_k_* will change under the influence of *v_l_*. Panel B shows the rate of change over time for *v_k_* as a result of *f_kl_* and *v_l_*. Note that the rate of change is offset from *v_k_* so that higher trait values will grow faster than lower trait values. Panel C shows how, over time, this offset will drive the trait-density distribution of the focal class *v_k_* to shift its mean in the direction of the gradient of *f_kl_*. Thus, at time *t* + Δ*t*, the mean trait *χ_k_* of *v_k_* will have shifted to become greater. **(D–F)** Causes and effects of term (*iv*) of Eqs. 5. Panel D shows trait-density distributions *v_k_*(*ξ,t*) and *v_l_*(*ξ,t*) of focal class *k* and source class *l* at time *t* along with a rate function *f_kl_* describing how *v_k_* will change under the influence of *v_l_*. Panel E shows the rate of change over time for *v_k_* as a result of *f_kl_* and *v_l_*. Note that the rate of change is offset from *v_k_* so that higher trait values will grow faster than lower trait values. Panel F shows how, over time, this offset will drive the trait-density distribution of the focal class *v_k_* to shift its mean in the direction towards the mean trait *χ_l_* of *v_l_*. Thus, at time *t* + Δ*t*, the mean trait *χ_k_* of *v_k_* will have shifted to become closer to *χ_l_*. **(G–I)** Causes and effects of term (*v*) of Eqs. 5. Panel G shows trait-density distributions *v_k_*(*ξ, t*) and *v_i_*(*ξ, t*) of focal class *k* and source class *l* at time *t* along with a rate function *f_kl_* describing how *v_k_* will change under the influence of *v_l_*. Panel H shows the rate of change over time for *v_k_* as a result of *f_kl_* and *v_l_*. Note that the rate of change is proportionally higher around the mean trait *χ_k_* of *v_k_* so that trait values closer to *χ_k_* grow faster than trait values further away. Panel I shows how, over time, this higher growth around *χ_k_* will drive the trait-density distribution of the focal class *v_k_* to shift its variance *V_k_* to become lower. Thus, at time *t* + Δ*t*, the standard deviation 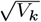 of *v_k_* will have shifted to become smaller. **(J–L)** Causes and effects of term (*vi*) of Eqs. 5. Panel J shows trait-density distributions *v_k_*(*ξ, t*) and *v_i_*(*ξ, t*) of focal class *k* and source class *l* at time *t* along with a rate function *f_kl_* describing how *v_k_* will change under the influence of *v_i_*. Panel K shows the rate of change over time for *v_k_* as a result of *f_kl_* and *v_i_*. Note that the rate of change is proportionally higher around the mean trait *χ_k_* of *v_k_* so that trait values closer to *χ_k_* grow faster than trait values further away. Panel L shows how, over time, this higher growth around *χ_k_* will drive the trait-density distribution of the focal class *v_k_* to shift its variance *V_k_* towards the variance *V_i_* of *v_i_*. Thus, at time *t* + Δ*t*, the standard deviation 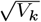 of *v_k_* will have shifted to become closer to the standard deviation the standard deviation 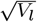 of *v_l_*. **(M–O)** Causes and effects of term (*viii*) of Eqs. 5. Panel M shows trait-density distributions *v_k_*(*ξ,t*) and *v_i_*(*ξ,t*) of focal class *k* and source class *l* at time *t* along with rate functions *f*_*kl*1_, *f*_*kl*2_, and *f*_*kl*3_, each describing one way of how *v_k_* will change under the influence of *v_l_*. Note that for *f*_*kl*1_, there is no directional selection, and only term *(viiia)* will play a role. Panel N shows the rate of change over time for *v_k_* as a result of each *f_kl_* and *v_l_*. The rate of change is offset from *v_k_* so that trait values at low density will grow faster than trait values at high density. Note also that the rate of change is more offset for *f*_*kl*2_ and less offset for *f*_*kl*3_ compared to *f*_*kl*1_. Panel O shows how, over time, these alterations to the offsets will drive the trait-density distribution of the focal class *v_k_* to shift its variance *V_k_*. This change is more pronounced for *f*_*kl*2_ and less for *f*_*kl*3_ compared to *f*_*kl*1_ due to the positive and negative gradients of *f*_*kl*2_ and *f*_*kl*3_ respectively. Thus, at time *t* + Δ*t*, the standard deviation 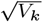 of *v_k_* will have shifted more for *f*_*kl*2_ and less for *f*_*kl*3_ compared to the shift caused by term (*viiia*).

Equation 5a describes the rate of change of the total density *uik* of species *i* in class *k*. Term (*i*), *per-capita growth,* describes the per-capita rate of process *m* between class *k* and class *l* evaluated at the mean-trait vector of species *i* in class *l* but is corrected by the a second-order term emanating from the Taylor expansion (Eq. 7a). This second-order correction is often referred to as the ‘genetic load’ in the quantitative genetics literature (Lande, 1979; Kirkpatrick and Barton, 1997), since if the species is distributed around a fitness optimum, the growth rate suffers a penalty due to the maladapted individuals around the optimum. The equations capture how the total densities of species change both due to within-class processes (*l* = *k*) such as local birth and death, and between-class processes (*l* ≠ *k*) such as dispersal between patches (c.f., Fig. 1A).

Equation 5b describes the rate of change of the mean-trait vector *χ_ik_* of species *i* in class *k*. Term (*ii*), *relative density weight,* weighs contributions by the relative total densities of classes *k* and *l*, so that if the density in the focal class *k* is much greater than that in the source class *l*, the ecological processes going from *l* to *k* will only have a marginal impact on the mean-trait vector in class *k*. Conversely, if the total density in class *l* is much greater than in class *k*, the ecological process going from *l* to *k* will have a large impact on the mean-trait vector in class *k*. The change of the mean-trait vector is then governed by two terms, (*iii*) and (*iv*). The first term, (*iii*), *directional selection,* describes the effect of directional selection in process *m* in class *l*, pushing *χ_ik_* in the direction of maximum increase of 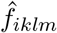, at a rate mediated by the available variance of species *i* in class *l*, *V_il_*. This effect comes about since the process in the source class will produce more trait-density on the side of the mean in which the slope is pointing, and less on the other side (Fig. 2A-C). The second term, (*iv*), *mean-trait flow,* describes how mean traits are homogenized by between-class transitions, where the rate of homogenization is governed by the per capita rate function 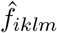, so that the mean trait of the focal class, *χ_ik_* will change in the direction of the mean trait in the source class, *χ_il_* (Fig. 2D-F). For within-class processes (*l* = *k*) we note that the relative density weight (*ii*) = 1 and the mean-trait flow (*iv*) = 0, meaning that for within-class processes only directional selection (*iii*) is relevant. If the process under consideration is trait-independent, such as for a constant dispersal rate between patches, the directional selection (*iii*) would be equal to zero, but mean-trait flow (*iv*) could still contribute towards changing the mean-trait vector in class *k* if the mean-trait vectors in *l* and *k* are different.

Equation 5c describes the rate of change of the variance-covariance matrix *V_ik_* of species *i* in class *k*. Like for the mean-trait vectors, the changes are weighted by the relative densities between class *k* and *l*, term (*ii*). The dynamics of the variance-covariance then depends on five terms. The first term, (*v*), describes *stabilizing/disruptive selection* resulting from the process with rate *f_iklm_*. Roughly speaking, if individuals close to the mean-trait vector *χ_il_* of species *i* in class *l* contribute more than individuals away from this optimum to process 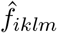 then the curvature as measured by 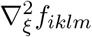 will be negative, which will contribute to a decrease in variances (Fig. 2G–I). Conversely, if individuals close to the mean contribute less, the curvature will be positive and this will contribute towards increases in variances. The second term, (*vi*), *variance-covariance flow,* homogenizes variance-covariances between classes, so that having trait-density flow from class *l* to *k* contributes towards driving the variance-covariance *V_ik_* of species *i* in class *k* closer to the variance-covariance *V_il_* in class *l* (Fig. 2J–L). The third term, (*vii*), *mutation,* is the contribution to variance-covariance from mutations in process *f_klm_*, which will contribute towards increasing variances and aligning the covariances of the species with the covariances of the mutational variance-covariance matrix *M_klm_*. Note that typically for most processes under consideration *M_klm_* would be zero, as in the two examples depicted in Fig. 1, where only birth processes are assumed to have mutations associated with them. How much mutations contribute towards variance increase also depends on the rate 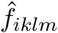. Thus, for example, in a system with high birth and death rates with mutations associated with births the mutations would have a stronger impact on variance-covariance than in a system with low birth and death rates even if net per capita growth were the same in both systems. Terms (*viii*), *effects of class-local adaptation on variance-covariance,* describe the effects of variability in mean-trait vectors between classes on the variance-covariance within each class. In the case where the classes are spatial patches, a species having different mean traits for different patches would simply be referred to as ‘local adaptation’, and we adapt this moniker here to the broader context of any class-structured population. The fourth term, *(viiia), between-to-within class variation*, describes how variances are increased and covariances altered by the differences in mean-trait vectors between classes, converting between-class variation into within-class variation (Fig. 2M–O). The final term, *(viiib), class-local adaptation and directional selection interaction,* comprises two terms which are the transposes of one another so that their sum is symmetric. They describe the interaction between directional selection and between-class differences in mean-trait vectors. This interaction is not trivial when the dimensionality is higher than one, but roughly speaking, variances will decrease when the mean trait difference and the selection gradient point in opposite directions, and increase if they are aligned (Fig. 2M–O). For within-class processes (*l* = *k*), the relative density weight (*ii*) will be equal to one, and only stabilizing/disruptive selection (*v*) and mutations (*vii*) will be nonzero, making these the only contributing factors. If the process *f_klm_* under consideration is trait independent, stabilizing/disruptive selection (*v*) and class-local adaptation and directional selection interaction (*viiib*) will be zero, but mutations (*vii*), variance-covariance flow (*vi*), and between-to-within class variation (*viiia*) can still contribute to changes in the variance-covariances.

In section B1.2 of Box 1, we showcase a concrete example of moment equations for the two-patch model.

#### Box 1: A two-patch model

In this box we showcase a simple example of our generic class-structured equations in the form of a two-patch model. This example model is used throughout the methods and models section to illustrate various facets of the theory.

##### Trait-space equations

For the two-patch model, we consider a system where consumers compete for two resources *R*_1_ and *R*_2_ and where the two classes are two patches, *A* and *B*. We assume that resources are renewed chemostatically on each patch, but that the supply of *R*_1_ is greater on patch *A* and that the supply of *R*_2_ is greater on patch *B*. We assume that the consumers consume resources with a type-I (linear) functional response, where the affinities *α*_1_(*ξ*) and *a*_2_(*ξ*) for *R*_1_ and *R*_2_ respectively are in a trade-off parameterized by a scalar trait *ξ* ∈ ℝ that is under selection. The trade-off is such that *ξ* = 0 yields a consumer with equal affinity for both resources, *ξ* > 0 implies a greater affinity for *R*_1_, and *ξ* < 0 implies a greater affinity for *R*_2_. We also assume that this trade-off is generalist-favoring (weak) so that *α*_1_(*ξ*) + *α*_2_(*ξ*) is maximized for *ξ* = 0.

In a well-mixed system on a single patch this would result in a single generalist consumer being evolutionarily stable (Levins, 1962; MacArthur and Levins, 1964; Rueffler et al., 2006; Wickman et al., 2019). Let *v_A_*(*ξ*) be the trait-density distribution of consumers on path *A* and *v_B_*(*ξ*) be the trait-density distribution on patch *B*. We then have the equations

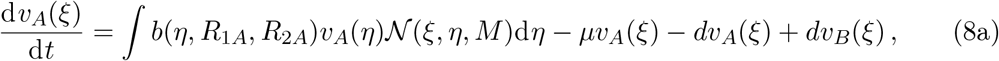

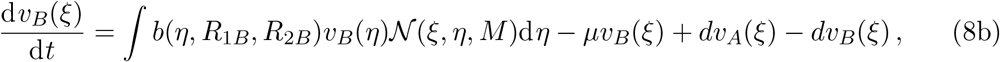

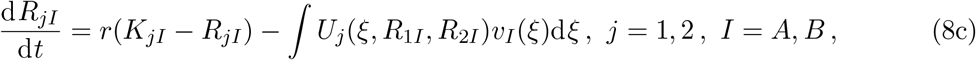

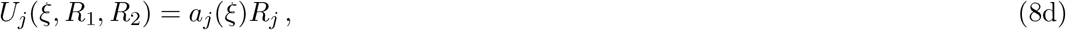

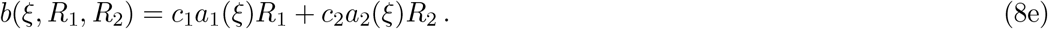

Here, *b* is the per capita birth rate, which depends on both the trait as well as the densities of resources *R*_1_ and *R*_2_ on a given patch. The consumers suffer a constant per-capita background mortality *μ*, and disperse between patches at a constant per capita rate *d*. Figure 1A provides an illustration of the system in the notation used for the generic equations (Eq. 2), where we can identify *f*_*AA*1_ = *b*(*ξ,R*_1*A*_,*R*_2*A*_), *f*_*AA*2_ = *b*(*ξ*, *R*_1*B*_,*R*_2*B*_), *f*_*AA*2_ = *f*_*BB*2_ = -*μ*, *f*_*AB*1_ = *f*_*BA*1_ = *d*, and *f*_*AA*3_ = *f*_*BB*3_ = -*d*. The birth rate has an associated mutational variance *M*, so that new individuals born from a parent with trait *η* are normally distributed around this trait with variance *M*, so that *M*_*AA*1_ = *M*_*BB*1_ = *M* in Fig. 1. The resource renewal rate for each resource on each patch is *r* and the maximal resource supply is *K_jI_* for resource *j* = 1, 2 on patch *I* = *A, B*.

##### Moment equations

By comparing the rate functions of the two-patch model (Eqs. 8) with the generic trait-space equations (Eq. 2), we can plug in our specific rate functions for the two-patch example into the generic moment equations (Eqs. 5) to get the moment equations for the two-patch system, which yields

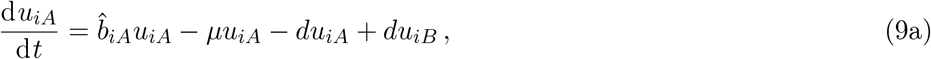

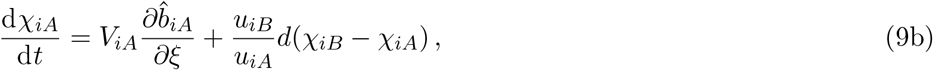

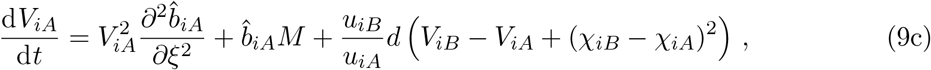

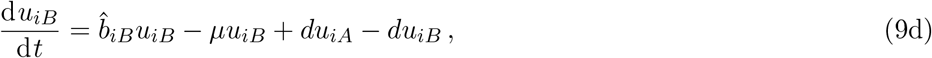

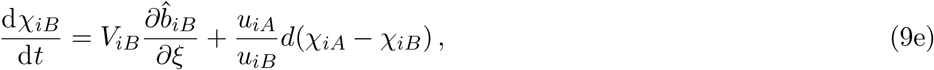

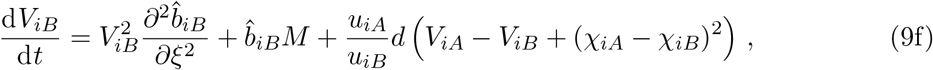

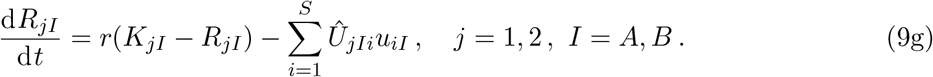

These equations are made less cumbersome by the fact that the only trait-dependent rates are the birth rates, which are within-class processes. As for the generic equations, we use a hat on a rate function to indicate that it represents a population rate or a Taylor approximation thereof. Thus, to Taylor order 3 we have e.g. that 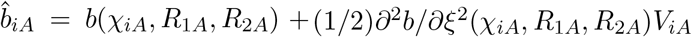. Equation 9a describes how the total density *u_iA_* for each species changes on patch *A* due to births, deaths, and dispersal to and from patch *B*. Equation 9b describes how the mean trait *χ_iA_* for each species changes on patch *A* due to directional selection on birth rates and mean-trait flow from patch *B*. Equation 9c describes how the variance *V_iA_* for each species changes on patch *A* due to stabilizing/disruptive selection in the birth rates, increases in variances due to mutations associated with the birth rate, variance-covariance flow from path *B*, and finally between-to-within class variation between the patches. Equations 9d–f describe the same processes on patch *B*.

We have also approximated the resource-uptake integral in Eq. 8c by Taylor-expanding the uptake function *U* around the mean-trait value of each species to second order. The details of how generic environmental variables such as resources can be incorporated into generic moment equations are given in Appendix A.

Figure 3 shows an example of numerical solutions of Eqs. 9 for two different rates of dispersal, *d*. For high rates of dispersal, the two-patch model is close to a well-mixed model and the eco-evolutionary equilibrium consists of a single generalist due to the generalist-favoring trade-off. For low rates of dispersal, a two-species equilibrium of two moderate specialists is attained due to the differential supply rate of the two resources on the two patches.

### Eco-evolutionary community assembly

In the two-patch model example (Box 1, Fig. 3) we saw that the model could yield either a one-species or two-species equilibrium, depending on the rate of dispersal between patches. A question that naturally arises then is: how can we tell how many species we need? For the relatively simple case of the two-patch model we could numerically solve the trait-space equations directly to determine this, but in more-complicated models the computational costs will in general be prohibitive. Thus, we would like some procedure for assembling an eco-evolutionary equilibrium that matches the equilibrium of the trait-space equations using only information from the moment equations. Here, we devise such a procedure by adapting two concepts from adaptive dynamics. The first criterion is local in trait space and determines when an evolutionary branching (Geritz et al., 1998) can take place. The second is a global invasion criterion to determine when an invader with traits that are not necessarily close to a resident can successfully invade the community. These conditions are then used to form so-called evolutionarily stable communities (Edwards et al., 2018).

**Figure 3:**
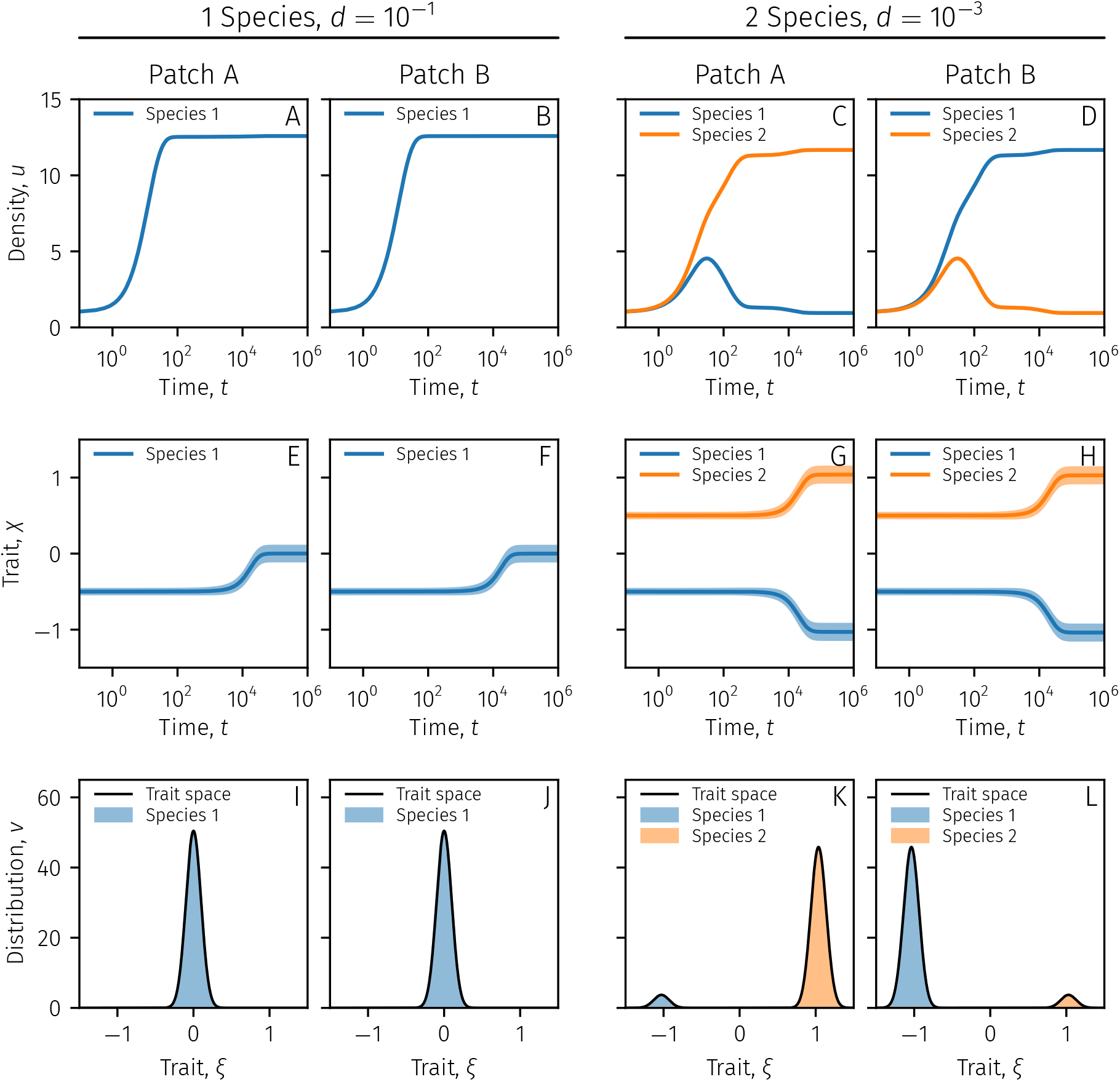
Illustration of numerical solutions of Eqs. 9 of the two-patch model (Box 1) for two parameter sets. The two left-side columns show solutions with one species for a high rate of dispersal (*d* = 10^-1^), and the two right-side columns show solutions with two species for a low rate of dispersal (*d* = 10^-3^). **(A–D)** Total densities over time on both patches for the two parameter sets. The densities change over time in accordance with Eqs. 9a,d. Note that for panels C–D each patch has one abundant species and one less abundant species, which is reversed between the patches. **(E–H)** Mean traits and standard deviations over time on both patches for the two parameter sets. The mean traits are shown as solid lines, and the colored area around each mean trait depicts one standard deviation around the mean. For high rates of dispersal, a single generalist evolves (Panels E–F), and for low rates of dispersal, two partial specialists evolve (Panels G–H). *χ*> 0 implies greater affinity for resource one and *χ* < 0 implies greater affinity for resource two. **(I–L)** Trait-density distributions at the final equilibrium on both patches for the two parameter sets. The numerical equilibrium is shown both for the numerical solution of the trait-space equations (Eqs. 8) as well as the species from the moment equations (Eqs. 9) depicted in the other panels. For these parameters, the moment equations approximate the equilibrium of the trait-space equations very well. **Parameter values**: *μ* = 0.1, *r* =1, *K*_1*A*_ = 1, *K*_1*B*_ = 0.4, *K*_2*A*_ = 0.4, *K*_2*B*_ = 1, *c*_1_ = 1, *C*_2_ = 1, *α* = 0.5, *M* = 5 · 10^-6^.

We note that our goal here is with respect to the ultimate assembled community in eco-evolutionary equilibrium. Our assembly procedure will in general not be a good approximation of the temporal dynamics of the trait-space equations leading up to the final community. We note also that our results here cannot be directly applied to questions of speciation, which would require the inclusion of genetic details not present in our framework.

### Evolutionary branchings

In adaptive dynamics, an evolutionary branching occurs when a species has evolved a trait in accordance with directional selection until directional selection ceases and the species finds itself at a fitness minimum causing the species to split in two (Geritz et al., 1998). This procedure cannot be directly translated to our moment equations due to the fact that we include trait variance and local adaptation. Instead, after Eqs. 5 have reached an equilibrium for *S* species we in turn split each species into two identical copies and examine the linear stability of the resulting *S* + 1 species equilibrium. If such a split-species equilibrium is unstable, we say that the system has undergone an evolutionary branching and keep the split-species pair. The details of the procedure is available in Appendix B, where we also present a concrete example for the two-patch model (Fig. B.1).

### Global invasion dynamics

Evolutionary branchings are not by themselves enough to ascertain global evolutionary stability across trait space as there may be invaders with substantially different traits from any resident species able to invade the community (Geritz et al., 1999; Klausmeier et al., 2020). Thus, once the moment equations for some fixed number of species (Eqs. 5) has reached equilibrium we would like to determine whether additional species that are not necessarily similar to any resident species can be added to the community. In adaptive dynamics, this is done by computing the invasion fitness of a invader with a fixed trait, assuming that the invader is so rare that it does not affect the environment it is invading. The invasion fitness is then defined to be the long-term exponential per capita growth rate of the invader, while the invader is still rare (Metz et al. 1992; see Caswell 2001 for structured populations).

Using this idea, we wish to construct a measure for our moment equations that tells us the ultimate fate of an introduced rare invader with some given trait. To do this, we also assume that the invader will remain so rare as to not affect the environment during the invasion process. We introduce an invader with a given mean-trait vector *χ*^inv^ that is the same in all classes, and with negligible variance-covariance, simulating the appearance of a single invading individual. We then let this invader mean-trait vector evolve until it has reached an equilibrium. We then let the variance-covariance of the invader evolve according to the moment equations until both the mean-trait vector and variance-covariance of the invader are in equilibrium. At this point, we can calculate the long-term exponential per capita growth rate of this invader. We will say that this is the invasion fitness of the invader. If the invasion fitness is positive, the invader will grow in the environment set by the resident community, and if the invasion fitness is negative or zero, it will not. We can thus associate an invasion fitness with any invading trait vector *χ*^inv^. The mathematical details of the procedure are available in Appendix B (Kremer and Klausmeier, 2013, used a similar approach in a non-structured model with fixed variance).

We have chosen to carry out the invasion process in steps, where first only the mean-trait vector is allowed to evolve, to avoid situations where the invader starts close to a fitness minimum in trait space, where the variance of the invader could potentially become too large before the invader has a chance to evolve a more favorable mean trait. The result is that each invading trait will trace a path in trait space until it reaches an equilibrium where its growth rate can be evaluated. However, calculating all such paths is not necessary for the community assembly process. If what we are interested in is to find any invader with positive growth rate, we can sidestep the first step where invaders evolve their mean-trait vectors by calculating the adaptive-dynamics fitness landscape. For rare invaders with no variance, evolving their mean traits then simply amounts to climbing this fitness landscape, and thus we need concern ourselves only with invading mutants with mean-trait vectors corresponding to the local peaks in the fitness landscape. Note that the adaptive-dynamics fitness landscape does not by itself yield sufficient information for our moment equations, as invaders may change their mean-trait vectors, and the population fitness of the invaders can change when variance-covariance is taken into account. An example of this phenomenon for the two-patch model is depicted in Fig. B.2 in Appendix B, where the adaptive dynamics fitness landscape indicates an evolutionary branching into two species, whereas the trait-space and moment equations admit only one.

### Assembly protocol

We can now use the moment equations, the branching condition, and the invasion process to assemble an eco-evolutionarily stable community. To assemble a globally eco-evolutionarily stable community we start with an arbitrarily specified community of *S* species in *K* classes with total densities 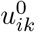, mean trait vectors 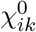 and variance-covariance matrices 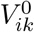. We then proceed along the following steps.

1. We let the community evolve according to Eqs. 5 until they reach equilibrium.
2. We check each species for evolutionary branching. In case of a branching we split the species undergoing branching into two new species and then return to step 1, letting the new community of *S* + 1 species evolve according to Eqs. 5 (as in e.g., Fig. B.1).
3. In case the moment equations for the resident community reach equilibrium and there are no branchings, we use our invasion scheme to see if any invader with positive invasion fitness exists. As described above, we first compute the adaptive-dynamics fitness landscape and locate its peaks. We then let an invader with a mean-trait vector corresponding to each peak invade in turn and let its mean-trait vectors and variance-covariances evolve until they all reach an equilibrium, after which we calculate the invasion fitness of each invader. If any invader with positive invasion fitness is found, the invader with maximum invasion fitness is added to the community with a small density, and we return to step 1, and let the new community evolve once again according to Eqs. 5.

We continue going through these steps until we have reached a community where Eqs. 5 are in equilibrium and no more invaders with positive invasion fitness can be found. This community is thus eco-evolutionarily stable. An illustration for this case is depicted for the two-patch model with asymmetric resource supplies in Fig. 4.

**Figure 4:**
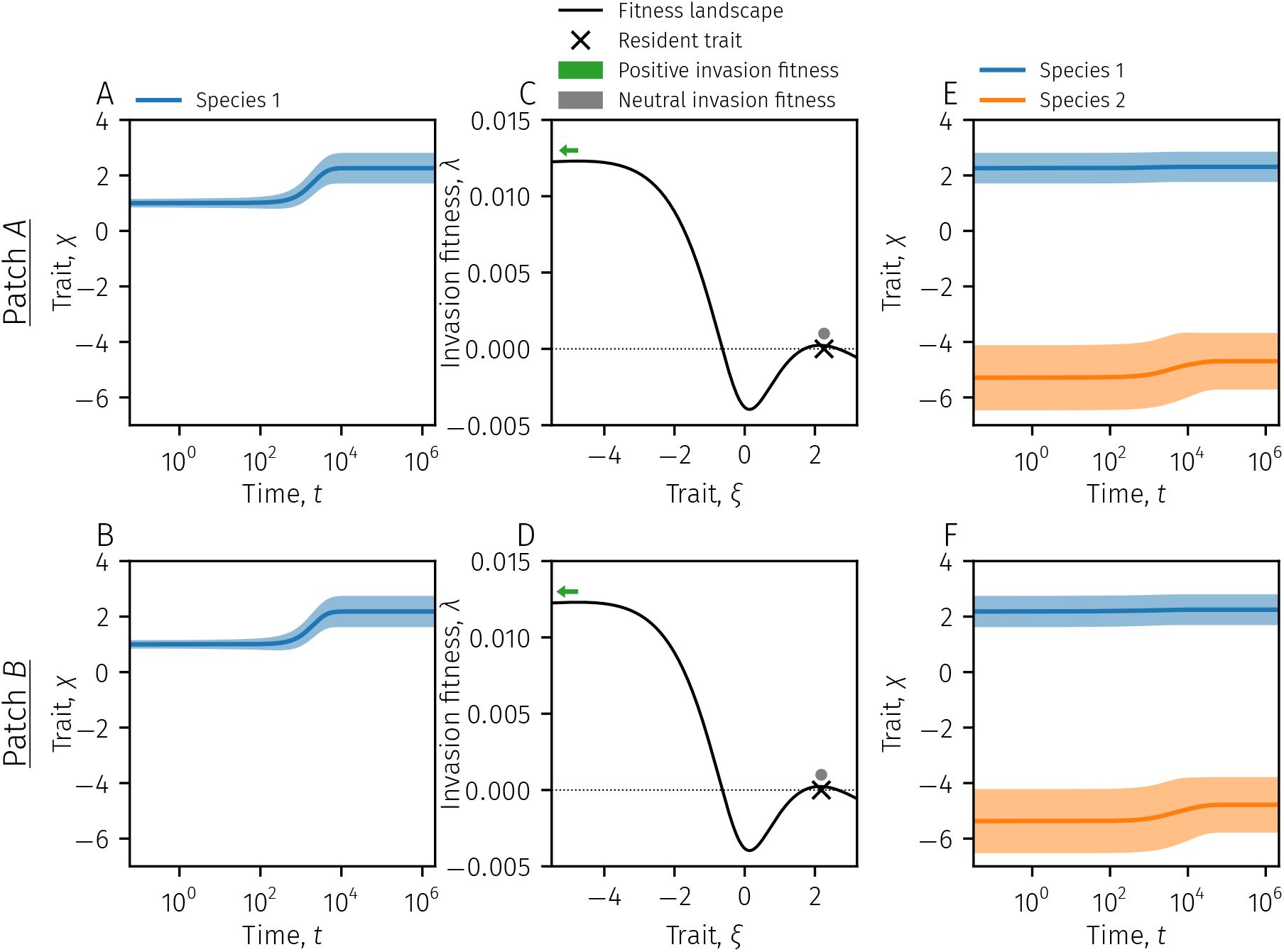
Illustration of the community-assembly procedure for the two-patch model (Box 1) with asymmetric resource supplies. **(A–B)** The species assembly process starts with one species, and the mean traits are evolved according to Eqs. 9 until equilibrium. **(C–D)** At equilibrium, the adaptive-dynamics fitness landscape is calculated, and an invader is tested at each local peak in the fitness landscape. Panels C and D show that one invader with *χ*^inv^ ≈ −4.7 (green arrows) and one with *χ*^inv^ ≈ 2.1 (gray dots) are tested. The bases of the green arrows show where the invader mean traits start, and the tip of the arrows show where they end up after their local mean traits and variances equilibrate. The invader with *χ*^inv^ ≈ 2.1 is close to the resident and ultimately ends up with the same mean trait and variance and resident on both patches, and hence neutral (zero) invasion fitness. While the invader close to the resident also adjusts its mean traits in response to trait variance, this movement is so slight it is not depicted. The invader with *χ*^inv^ ≈ −4.7 however ends up with positive invasion fitness and will be added to the community. **(E–F)** The community is then once again evolved according to Eqs. 8 until the community is once again at equilibrium. At this point, no more invaders with positive invasion fitness can be found (not shown). **Parameter values**: *μ* = 0.1, *r* = 1, *K*_1*A*_ = 10, *K*_1*B*_ = 0.1, *K*_2*A*_ = 1, *K*_2*B*_ = 1, *c*_1_ = 1, *c*_2_ = 1, *α* = 0.4, *M* = 10^-3^, and *d* = 10^-2^.

Following this process allows us to find an eco-evolutionarily stable community with the correct number of species in the final community while including trait variance-covariance. This allows us to explore parameter space to determine how different environments and non-evolving species properties may generate different eco-evolutionarily stable communities. In Fig. 5 we show an example for the two-patch model where we assemble eco-evolutionarily stable communities for different dispersal rates *d* and mutation variances *M*. As surmised, for high rates of dispersal a single generalist makes up the community, whereas low rates of dispersal engender a two-species community of moderate resource specialists.

**Figure 5:**
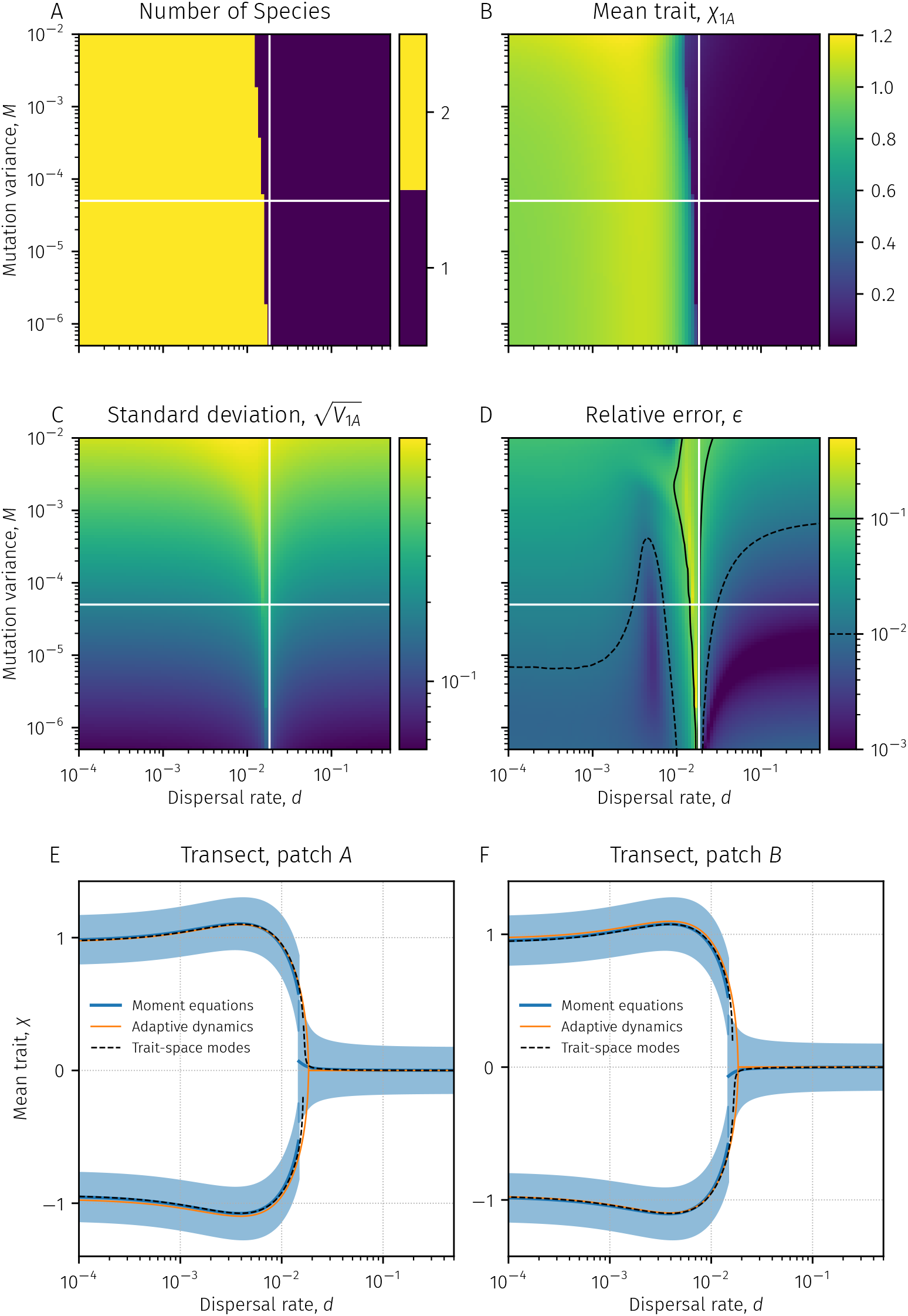
Properties of eco-evolutionarily stable communities in the two-patch model (Box 1) for various rates of dispersal and mutation variances. **(A)** Number of species in the assembled evo-evolutionarily stable community. The vertical white line indicates the dispersal rate for which the adaptive-dynamics version (zero variance) of the two-patch model undergoes an evolutionary branching from one to two species. The horizontal white line indicates the transect displayed in Panels E and F. **(B)** Mean-trait value of the resource one specialist on patch *A*, *χ*_1*A*_. **(C)** Standard deviation of trait distribution of the resource one specialist on patch *A*, 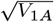. **(D)** Relative error *ϵ* between the moment equations and trait-space equations at the eco-evolutionarily stable state. The error is calculated as 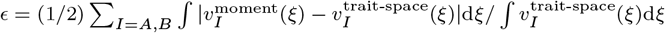. Thus, perfect agreement between the distributions would mean that *ϵ* = 0. For reference, the relative error between moment and trait-space solutions depicted in Fig. 3K–L is *ϵ* ≈ 8.97 · 10^-3^. **(E–F)** Fuller display of the transect depicted as a horizontal white line in Panels A–D on patches *A* and *B* respectively. The mean traits of the species are depicted as blue lines, and one standard deviation is depicted around these mean traits as blue areas. The adaptive-dynamics eco-evolutionarily stable communities are depicted as orange lines. The modes (locations of local maxima) of the trait-space solutions are depicted as broken black lines. For all panels, parameters other than *M* and *d* are as in Fig. 3.

In Fig. 5E-F we also display eco-evolutionarily stable communities in the adaptive-dynamics sense, which can be considered the limiting behavior of the moment equations when there is no trait variance. Apart from the obvious difference that our moment equations track the variance of each species, the moment-equation eco-evolutionarily stable communities show two notable differences. The first is that non-zero variance slightly delays the transition to a two-species community compared to adaptive dynamics. This is likely due to both the intraspecific variation itself as well as the local adaptation—that the mean traits can differ from patch *A* and *B* for the same species—it engenders. The second difference is the local adaptation itself for the sink-population species on the off-patch for each species. Due to the intraspecific variation, each species is slightly more generalist on the patch on which it is least adapted, as opposed to the adaptive-dynamics case where a single trait describes the entire species in all classes.

The price we pay for using the moment equations in our assembly procedure is that the accuracy of the moment equations in representing the trait-space equations can be variable (Fig. 5D). For the cases we have tested, the moment equation solutions at eco-evolutionary equilibrium typically have good quantitative accuracy, nearly always good qualitative accuracy and only rarely outright disagreement with the trait-space solutions. When there is approximation inaccuracy, it is primarily driven by two things. First, our assembly framework performs worse when the number of species in the trait-space solutions is ambiguous, in this example between one and two species. This is likely due to our assumption that each species can be treated as if it were in reproductive isolation. However, around the transition point the trait-space equations tend to converge on trait-density distributions that are multimodal but without well-separated peaks; for this case, the notion of splitting this distribution into two reproductively isolated distributions does not always work well. Second, under high mutation variances, the parameter space generating this ambiguity is increased. The assembled moment solutions can however be surprisingly effective even when mutation variances—and hence the variances of the species—are high, as seen towards the top left of Fig. 5D. In Fig. C.1 (Appendix C) we show two sample solutions for intermediate and high mutation variances. For the larger mutation variances, approximately *M* ≥ 3 · 10^-3^, and low dispersal rates the model undergoes a regime shift as the trait variation of the species become comparable to the distance between the individual patch optima *χ_A,opt_* ≈ 1 and *χ_B,opt_* ≈ –1, and the trait-space equations yields unimodal distributions on both patches even for low rates of dispersal. Our assembly model yields two species, but the mean traits of the species are close enough and the variances large enough that the sum of the normal distributions is unimodal (Fig. C.1).

In addition to this, for some small slivers of parameter space, the moment equations do not converge on a stable community, but are instead characterized by eco-evolutionary cycling (Fig. C.2). For Fig. 5 the resolution in parameter is coarse-grained enough that we do not hit any of these points. Experiments with using second-order Taylor expansions did yield larger areas in parameter space characterized by cycling (data not shown) compared to the third-order expansions (Eqs. 7) used to generate Fig. 5.

In spite of these shortcomings, the moment equations coupled with our assembly procedure works remarkably well outside of the small band of parameter space where the number of species is ambiguous (Fig. 5D).

## Additional example: A stage-structured resource-competition model

To illustrate the application and utility of our methods, we present here an additional example of eco-evolutionary community assembly taking intraspecific variation into account. The two-patch model we used to illustrate the generic equations was deliberately simple. First, it involved only a single trait. Second, the transition between classes (dispersal) was constant, and did not depend on traits. However, such simplifications cannot always be made. In stage-structured models where juveniles mature and adults give birth to new juveniles, these rates will often depend on the traits of the species. To demonstrate such a case, we here adapt a model by De Roos et al. (2007) of a population subdivided into two stages, juveniles with trait-density distribution *v_J_* and adults with trait-density distribution *v_A_*, competing for two resources *R*_1_ and *R*_2_. Both the maturation rate and the birth rate depend on how much resources the two stages acquire, as well as the traits associated with resource acquisition.

The trait-space equations read

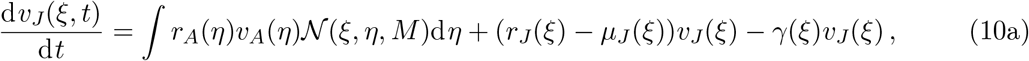

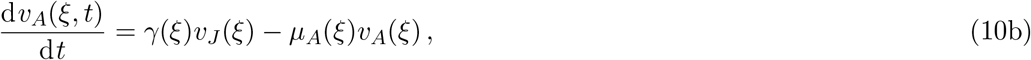

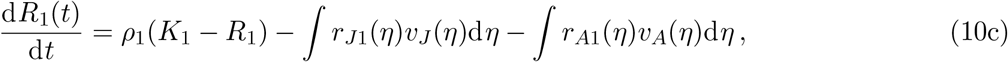

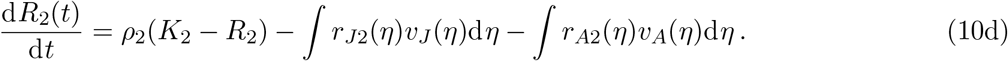

Here, the distributions *v_J_*(*ξ*) and *v_A_*(*ξ*) characterize the distributions of biomass density in trait space for juveniles and adults respectively. Both adults and juveniles consume the substitutable resources *R*_1_ and *R*_2_ with a type-II functional response at rates *r_Ij_* for *I* = *A, J*, and *j* = 1, 2. A conceptual illustration of the trait-space equations is depicted in Fig. 1B.

The adults use all of their intake of resources *R*_1_ and *R*_2_ to give birth to juveniles at a per capita rate of *r_A_* = *r*_*A*1_ + *r*_*A*2_, and suffer a mortality at a per capita rate of *μ_A_* Juveniles grow at a per capita rate *r_J_* = *r*_*J*1_ + *r*_*J*2_, suffer a mortality rate of *μ_J_*, and transition into adults at a rate *γ* that depends both on juvenile growth and mortality (De Roos et al., 2007). The specific forms of these functions are given in Appendix C. In the absence of consumers, the resources are renewed chemostatically at rates *ρ_j_* up to maximal supplies of *K_j_* for resource *j*.

We let the trait vector *ξ* = (*ξ*_1_, *ξ*_2_) ∈ ℝ^2^. The first trait *ξ*_1_ parameterizes trade-offs in resource affinities, so that juveniles experience a trade-off between their affinities *a*_*J*1_ and *a*_*J*2_ for *R*_1_ and *R*_2_ respectively, and adults experience a trade-off between their affinities *a*_*A*1_ and *a*_*A*2_. While the same trait component 1 parameterizes the trade off in resource affinity for both juveniles and adults, we let the shapes differ so that the juveniles experience a specialist-favoring trade-off while adults experience a generalist-favoring trade-off. Such differences could come about as foraging behavior changes between life stages. The second trait *ξ*_2_ parameterizes a trade-off between juvenile mortality *μ_J_* and adult mortality *μ_A_*. Such trade-offs between juvenile and adult mortality could come about through, e.g., parental investment or risk taking on behalf of juveniles. The parameter values used in our numerical solutions can be found in Appendix C.

Following our general procedure, to get our moment equations we now assume that the solutions to Eqs. 10a–b are composed of a set of *S* discrete peaks for both juveniles and adults, and approximate the solutions by

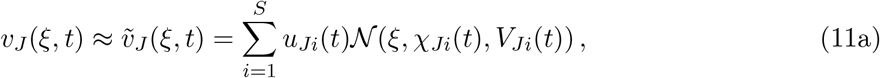

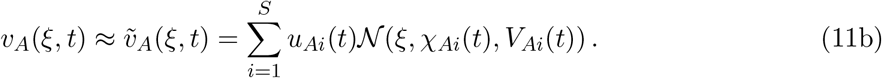

This means that each species *i* = 1,2,…,*S* is represented by six variables: the total densities of juveniles, *u_Ji_*, and adults, *u_Ai_* the mean trait vector for juveniles, *χ_Ji_*, and adults, *χ_Ai_* and the trait variance-covariance matrices for juveniles, *V_Ji_*, and adults, *V_Ai_*. We can derive the moment equations for these quantities by plugging in the expressions for the trait-space equations for this model into our derived generic equations (Eq. 5) yielding the moment equations for this stage-structured model that read

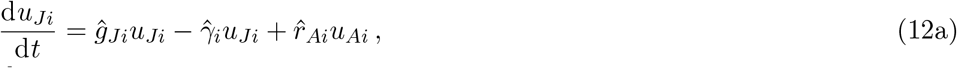

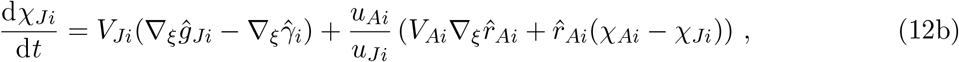

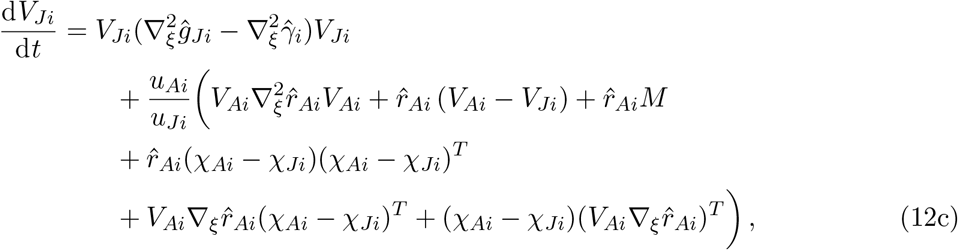

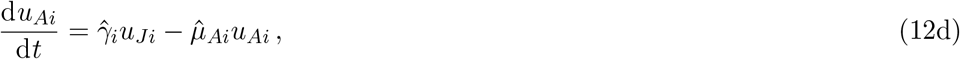

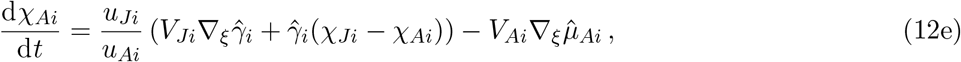

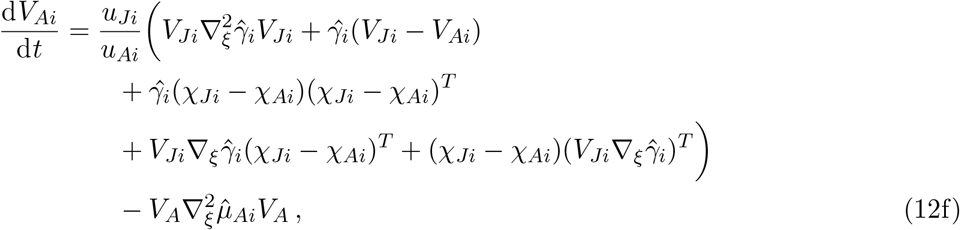

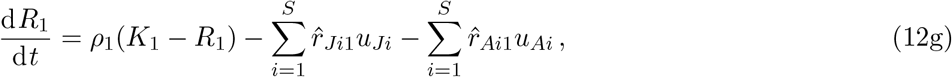

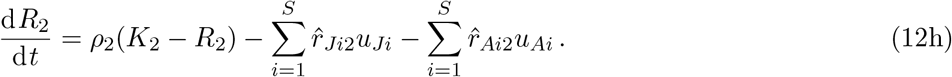

While somewhat involved, the various terms admit to ecological interpretation, and can be understood in the following way. First, recall that we use functions with a hat as a shorthand for the third-order Taylor approximation of the population rates, see Eqs. 7. We also let *g_Ji_* be the net per capita growth of juveniles, i.e., *g_Ji_* = *r_Ji_* – *μ_Ji_*. For the purposes of readability we have also suppressed any dependence on function arguments in the equations above.

The terms of Eq. 12a describe in turn net growth in mass of juveniles, the removal of juveniles that transition into adults, and the addition of mass due to birth by adults. While suppressed for the sake of notation clarity, these quantities depend also on the variance-covariance matrices of the juveniles and adults through the third-order Taylor approximation (Eqs. 7).

The changes to the mean trait vector of the juveniles *χ_Ji_* (Eq. 12b) are divided into two terms. The first, with a matrix factor *V_Ji_*, describes how directional selection on the juvenile processes of net growth and transition to adulthood moves the mean trait of the juveniles. This is because juveniles with certain traits will have higher net growth than others, and this will push the mean trait of juveniles in the direction of the gradient 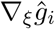. Conversely, some juveniles with certain traits will mature into adulthood more quickly than others, which removes them from the juveniles and thus moves the mean trait in the opposite direction of the gradient 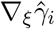. The strength of both these effects depends on the how much variation *V_Ji_* there is for a given species *i*. The second term, with a scalar factor *u_Ai_*/*u_Ji_* describes the influence of births from the adults on the mean trait of the juveniles. This is, in turn, composed of two terms. The first term, 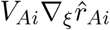, describes how directional selection in the birth rate of juveniles by adults contributes towards change of juvenile mean traits. This effect comes about since adults with different traits give birth at different rates, and more offspring will be produced along the direction of the gradient 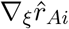. This effect is stronger if the variation among the adults *V_Ai_* is larger. The second term, 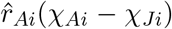, describes how the juvenile mean trait is pushed in the direction of the adult mean trait, homogenizing juvenile and adult mean traits. This is due to the fact that adults of species *i* will on average give birth to juveniles with trait *χ_iA_*, and so, if there is a difference between juvenile and adult mean trait, the juvenile trait will become more similar to the adult mean trait over time. The overall factor *u_Ai_*/*u_Ji_* comes about through the fact that how much effect birth of juveniles by adults has on the juvenile mean trait depends on the mass ratio between the two stages. If there are few adult compared to juveniles, then the birth of new juveniles will have little effect on the mean trait of all the juveniles.

Equation 12c describes the changes to the variance-covariance matrix of the juveniles over time. The first term, 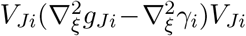, describes the effects of stabilizing or disruptive selection of the net growth and transitions. Thus, for example, if the curvature of the net growth function, 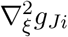, is negative around a mean trait component, this will reduce the variance of that trait component over time. There are then several terms with a factor *u_Ai_/u_Ji_*. The first term, 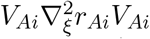, reflects the effects of stabilizing or disruptive selection in the birth rates. The second term, 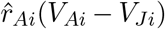, describes the homogenization of variance-covariance matrices between the stages, so that this term changes the juvenile variance-covariance to be more similar to that of the adults over time. The third term, 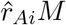, describes the increase to juvenile trait variances due to mutations. Next, the term 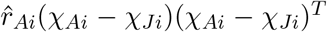 describes how variances are increased, and covariances changed over time due to differences in means between the juveniles and adults. This reflects that if the means of the adults are away from the means of the juveniles, then birthed juveniles will be away from the mean of the extant juveniles, and thus increase variance. The next two terms, 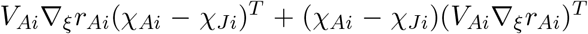, which are the transposes of one another, take into account the interaction of distance between the means of the stages and effects of directional selection in the birth rates.

The terms in the equations for the adults are mostly analogous to those for the juveniles, with the difference that the maturation function *γ* now plays the role that the birth rate function *rA* did for the juveniles, as this is the function that determines the rate of transition from the juvenile to the adult stage. Additionally, as we can see in the trait-space equations (Eqs. 10), we assume that a juvenile with trait matures into an adult with the same trait *ξ*, and thus no mutation term enters into the equations for adult variance-covariance. Finally, we approximate the integrals in trait space in the resource equations (Eqs. 10c–d) to second order around the mean traits (see Appendix C for details).

Using these equations we let the population go through our evolutionary assembly process (Fig. 6A). The traits *ξ*_1_ and *ξ*_2_ parameterize the trade-offs in resource affinities and mortalities respectively. However, as the traits that actually matter for the ecology are the affinities and mortalities themselves, we depict this assembly process in the derived trait space of (*a*_*I*1_ – *a*_*I*2_) × (*μ_J_* — *μ_A_*) for *I* = *J, A* to get a better understanding of how the expressed traits that impact the ecology change. The final community of one moderate *R*_1_ specialist and one moderate *R*_2_ specialist comes about through the conflicting trade-offs for juveniles and adults. The specialist-favoring trade-off in resource affinity for the juveniles engenders selective pressure towards specialization into two species, but the generalist-favoring trade-off for the adults will eventually counter balance this disruptive force, and the community will settle on substantial, but not complete, specialization for both species. The differences between juveniles and adults in their resource affinities seen in Fig. 6A stem mostly from their their different trade-offs; the mean traits *χ_j_* and *χ_A_* differ little within the same species. The distributions in *ξ*_1_ × *ξ*_2_ space can be seen in Fig. C.4, where we also compare the numerical moment solutions for the eco-evolutionarily stable two-species community to numerical solutions of the trait-space equations.

**Figure 6:**
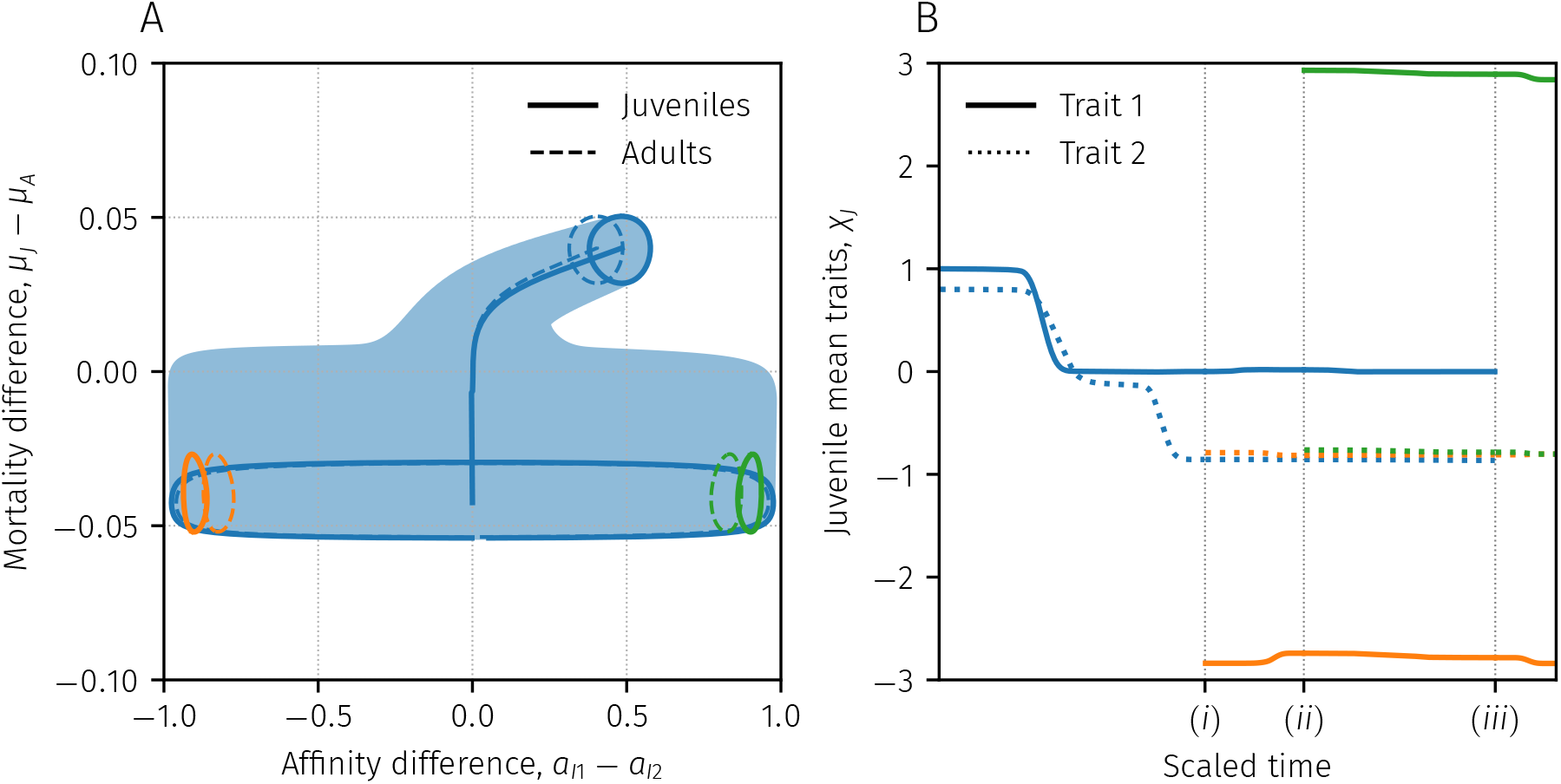
Community assembly in the stage-structured example. **(A)** Depiction of the assembly process and final eco-evolutionary equilibrium in the derived trait space (*a*_*I*1_ – *a*_*I*2_) × (*μ_J_* — *μ_A_*) for *I* = *J,A*. The eco-evolutionary process is started towards the top-right corner, where the evolutionary paths of the mean derived traits *a*_*I*1_ — *a*_*I*2_ and *μ_J_* — *μ_A_* are depicted as a solid blue line for juveniles and broken blue line for adults. The 95% transformed confidence ellipse for the juveniles is depicted at each time-point around the means in lighter color. The evolutionary process continues until the system reaches a one-species equilibrium. The 95% transformed confidence ellipses are shown as a solid blue ellipse (juvenile) and dashed blue ellipse (adult). This one-species equilibrium is subsequently invaded by two specialist species, and the final two-species equilibrium is depicted in orange and green. **(B)** Evolution and assembly of juvenile mean traits over time. Trait 1 (solid line) governs the resource-affinity trade-off and trait 2 (dotted line) governs the juvenile–adult mortality trade-off. The process starts with one species reaching an eco-evolutionary equilibrium, but is invaded by species two (orange) at (*i*). The two-species community subsequently reaches an eco-evolutionary equilibrium, which is in turn invaded by a third species (green) at (*ii*). With two specialists to compete with, the generalist species (blue) cannot persist and goes extinct at (*iii*). The new two-species community then evolves to a new equilibrium which is stable to further invasion, and the assembly process is complete. Time has been rescaled non-uniformly to illustrate the process.

## Discussion

In this paper we have presented a general framework for eco-evolutionary community assembly for class-structured communities that incorporates intraspecific trait variation. We have done so by deriving moment equations for the total density, mean trait, and trait variance covariance for each species in a community, and combining it with a procedure for determining whether additional species need to be added to a community in order for it to be closed to further invasion. Through examples, we demonstrated the application of the framework in a spatially-structure two-patch model and a stage-structured model with one juvenile and one adult stage.

Our framework builds on, and connects to, several other strands of eco-evolutionary theory. Considering a single class, and setting all mutations to zero, our framework reduces to the community-ecology framework of Wirtz and Eckhardt (1996) and Norberg et al. (2001), where the moments of the trait-distribution of an ensemble of species is tracked. This approach often includes an immigration term from a fixed species pool to maintain variance in the community. Such an immigration term could easily be incorporated into our framework by designating one of the classes a “species-pool class” and letting the internal rates of this class all be zero to keep the species pool fixed. Note that our formulation also includes cases where the species pool too consists of structured populations, so that, for example, juveniles and adults in a stage-structured model could immigrate at different rates from the species pool.

Although the assumptions going into the model at the outset are different, the moment equations derived for our model are also very similar to the equations derived under the assumptions of quantitative genetics. Assuming a single class, no mutations, and that phenotypic variation equals genetic variation (no environmental variation), our equations mirror those of Barabás et al. (2020), who derived equations for mean traits and variance-covariance under the assumptions of quantitative genetics for multiple traits. These similarities between trait-space approaches and quantitative genetics have been noted before (e.g., Débarre et al., 2013) and the two approaches have complementary strengths and weaknesses. Most notably, the assumptions of normality are less ad hoc in quantitative genetics, and each species is, as described by a normal distribution, well defined. Similarly, the reproductive isolation of species, which in our moment equations is an approximation, is in quantitative genetics based on the biological species concept. This, however, comes at the cost that diversification into multiple peaks cannot be easily incorporated as opposed to our approach here.

Finally, our moment equations closely resemble those derived in trait-diffusion approaches (Merico et al., 2014; Le gland et al., 2020), where mutations are generated by a diffusion process in trait space. Merico et al. (2014) derived the moment equations for well-mixed, single-species, single-trait populations and Le gland et al. (2020) extended this to multiple traits and spatial structure by way of reaction-diffusion equations in continuous space. For mutation kernels with small variance-covariance matrices without covariances, our mutation convolution integral is well approximated by such trait-diffusion processes (Kimura, 1965; Débarre et al., 2013), and our assembly framework is easily adapted to this setting, equipping the trait-diffusion approaches with a way of assembling eco-evolutionarily stable communities of several species.

Our framework for eco-evolutionary community assembly builds on key ideas from other frameworks where the ecological dynamics can be fruitfully described by differential equations and thus inherits some of their key strengths, but also some of their weaknesses. On the one hand, as opposed to adaptive dynamics (Metz et al., 1992; Dieckmann and Law, 1996; Geritz et al., 1998; Dercole and Rinaldi, 2008; Kremer and Klausmeier, 2013) where every species is entirely monomorphic with a single trait value, our framework incorporates intraspecific variation while still allowing us to perform an invasion analysis to assemble eco-evolutionarily stable communities. Our evolutionary branching criterion closely mirrors that found in adaptive dyanimcs. However, the global assembly process we have devised is a heuristic as opposed to the situation in adaptive dynamics where, at least in theory, the fitness landscape can always be computed to ascertain that a community is truly closed to further invasions. In practice, as evidenced by the examples we have presented here, our heuristic has proven reliable for community assembly for many scenarios, often covering most of parameter space, at least for modest mutation variances (Fig. 5D). Our approach is thus a good choice when intraspecific variation needs to be taken into account and the assembly of an eco-evolutionarily stable community is desired.

On the other hand, our framework takes into account replicative infidelity (mutations) as opposed to most moment-based frameworks (e.g., Norberg et al., 2001; Barabás and D’Andrea, 2016), which is a key part for our community assembly process. However, this too comes at cost, as evaluating the mutation integrals explicitly required us to assume that both species’ trait distributions and mutation kernels were normally distributed. In other moment-based frameworks, other so-called moment closures are also possible, which could be more appropriate depending on the problem under study. For example, Klauschies et al. (2018) and Cropp and Norbury (2021) used beta distributions as the approximating distribution to close their moment equations. Assumptions other than normality are likely possible while still retaining the core features of our approach, but each such differing assumption would require the re-derivation of nearly all moment equations. While a shortcoming, our comparisons between the trait-space equations and moment equations indicate that as long as normality in the distributions of birthed phenotypes from a parent phenotype is assumed, then our additional assumption that species’ trait distributions are normal seem not to affect the accuracy of the moment approximations by any large degree, as our eco-evolutionarily stable communities assembled by moment equations agreed very well with the corresponding traitspace equations. We note that although we assume normality in each species, the community trait-density distribution is the sum of these normals, which makes the community distribution much more flexible, including both multimodal distributions as well as highly non-normal uni-modal distributions (see Fig. C.1C-D for an example). The primary exception to good accuracy in the moment equations is when the number of species is ambiguous. In these instances the traitspace equations would yield solutions that could not easily be approximated by a sum of normal distributions that were assumed to be reproductively isolated.

While these drawbacks should be kept in mind, we nevertheless believe that our framework makes substantial progress in eco-evolutionary modeling with intraspecific trait variation. In a phytoplankton model Peeters and Straile (2018) compared trait-space equations without mutations to single-species moment equations, and concluded that single-species moment equations failed to provide any useful information when considering parts of parameter space where the trait-space equations diverged into multiple species. In a similar model using the trait-diffusion approach Le gland et al. (2020) noted that their trait-space equations sometimes exhibited multi-modality and speculated on the utility of modeling multiple modes, making the selection of how many modes to include based on functional groups. While multi-species moment models are not new (Sasaki and Dieckmann, 2011; Norberg et al., 2012; Barabás and D’Andrea, 2016), our assembly approach obviates the need for a-priori decisions on how many modes or species to include by assembling the correct number of species, at least as long as the number of species is not ambiguous.

One important benefit of incorporating intraspecific variation into an eco-evolutionary community assembly framework is that it makes possible to study a wider array of diversity-related questions. This allows theoretical explorations of e.g., the effects of productivity on diversity (Adler et al., 2011; Cusens et al., 2012) and the effects of diversity on ecosystem services (Cardinale et al., 2011) from a trait-based perspective while recognizing the importance of eco-evolutionariliy stable communities (Edwards et al., 2018). In recent years, there has been much interest in the effects of intraspecific variability (Albert et al., 2010; Violle et al., 2012) and many different metrics have been proposed and used (Schleuter et al., 2010; Fontana et al., 2016). Our class-based framework tracks not only the intraspecific trait variation, but also the difference in mean traits between species, as well as the difference in trait distribution between classes within each species. For the case when classes are spatial patches, this opens up the potential to study both how intraspecific and interspecific diversity, as well as alpha and beta diversity are shaped by conditions and influences outcomes.

Taken together we believe that our framework of unifying the community-assembly techniques of adaptive dynamics with the moment-equation approach to including intraspecific trait variation could be of great use to theoreticians and modelers seeking to take advantage of facets of both eco-evolutionary modeling frameworks. We also believe that being able to assemble eco-evolutionarily stable communities that accounts for trait variation could help address many ecological questions where intraspecific trait variation is thought to be important.

## Acknowledgments

This project was funded by NSF grant DEB-1754250 to E. Litchman and CAK. We thank Florence Dáebarre, Sáebastien Lion, Elena Litchman, and Bob Week for their comments on the work.

## Appendix A: Derivation of moment equations

### Calculating the time derivatives and general notation

#### Time derivatives of the moments

Let the trait-density distribution of a community be given by *v*(*ξ*) where *ξ* ∈ ℝ^*n*^ is a trait vector. Let *ψ* and *ω* index the trait components of *ξ* and let dots over symbols denote time derivatives. We now wish to consider the following moments of the trait-density distribution *v*:

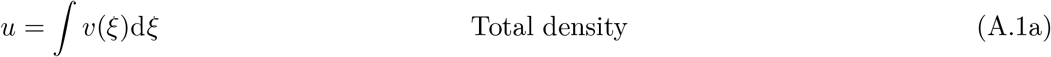

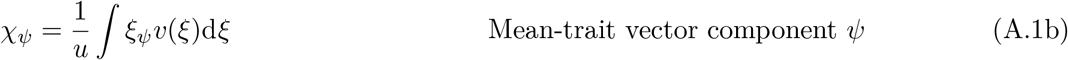

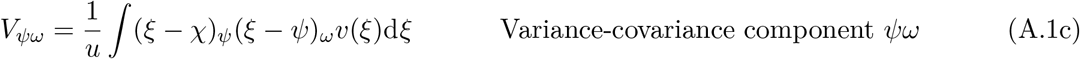

All integrals are over the entirety of ℝ^*n*^. Calculating the moment time derivatives then yields:

For the total density *u*:

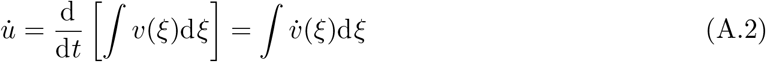

For the mean trait component *χ_ψ_*:

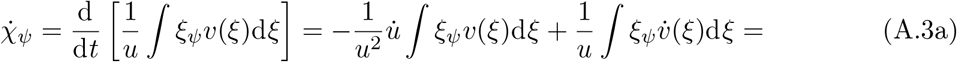

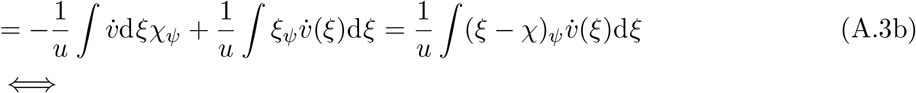

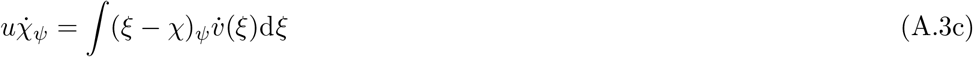

For the variance-covariance component *V_ψω_*:

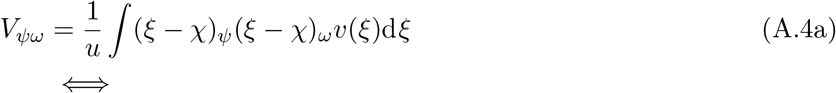

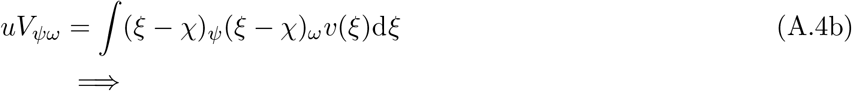

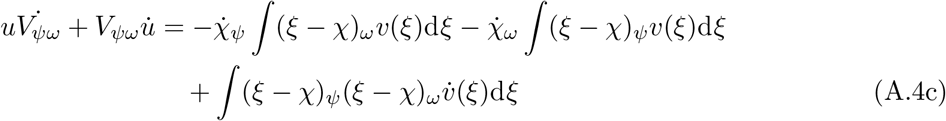

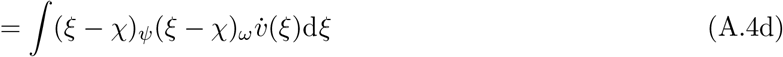

Together we thus have

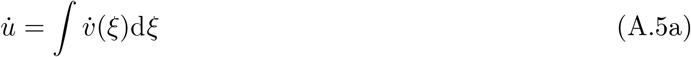

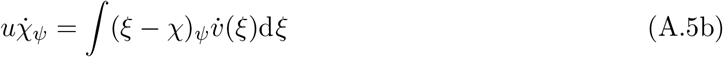

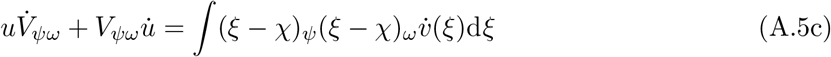

This way of expressing the relationships between the time derivatives of the moments and the time derivative of the trait distribution makes it easier to derive the moment equations for the various models, as we can plug in the right-hand side of any equation in place of 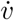 in Eqs. A.5.

#### A note on notation

The calculations for deriving the moment equations are not difficult in the sense that they require any complicated mathematical concepts or techniques. They are however rather complex in terms of notation and book keeping. To alleviate this problem, we introduce some useful notation.

First, we frequently employ so-called Einstein summation notation. This means that, unless otherwise indicated, repeated indices are summed over, so that for example

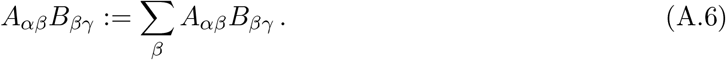

Second, we let indices on scalar quantities denote components of gradients, i.e., if *f* is a function from ℝ^*n*^ to ℝ, then

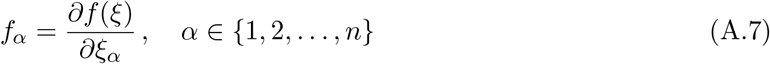

Thus, for example, in index notation we would write ∇_*ξg*_ as *g_α_*. Combining these two notational conventions, we can for example write the matrix-vector product *V*∇_*ξg*_(*χ*) as *V*∇_*ψαgα*_ and the matrix-matrix-matrix product 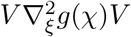 as *V*_*ψαgαβ*_*V_βω_*.

We will use Greek letters to indicate trait components. For these indices the Einstein summation convention will always apply. We will use Latin indices *i* and *j* for species, and *k* and *l* for classes, and finally *m* for processes. The Einstein summation convention will not apply to these indices.

### Deriving the moment equations

To make it easier to derive the moment equations for the trait space equations we will first set out to derive some intermediate results.

First, let 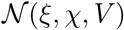 be a normal distribution probability-density function with argument *ξ*, mean vector *χ*, and variance-covariance matrix *V*. We can then calculate the first and second derivatives of this normal distribution with respect to its argument components:

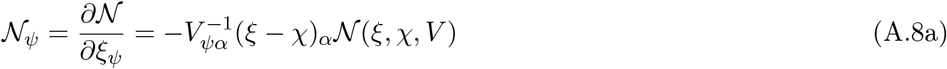

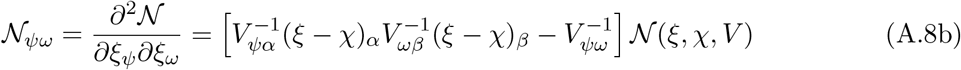

Furthermore, if *f* (*η*) is the rate of an ecological process for an individual with trait *η*, let the population level per capita rate for a population with mean-trait vector *ξ* and variance-covariance matrix *W* be given by

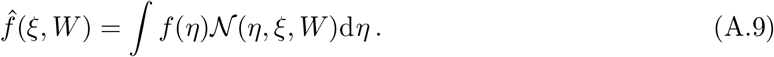

We can now differentiate the population-level rate with respect to the mean-trait components:

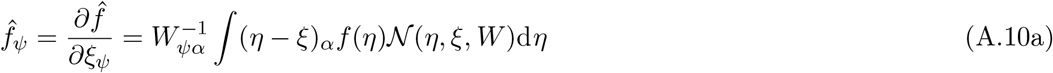

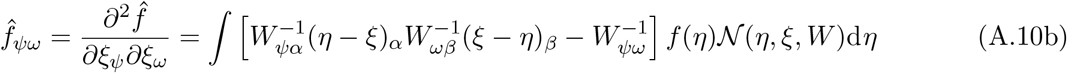

Using these results we can calculate two integrals that will appear later in our derivation:

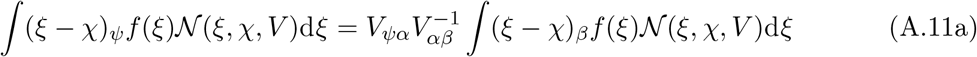

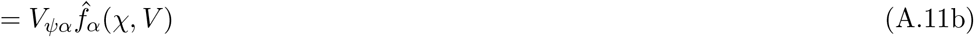

and

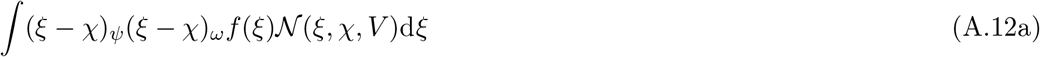

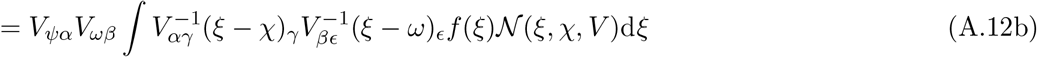

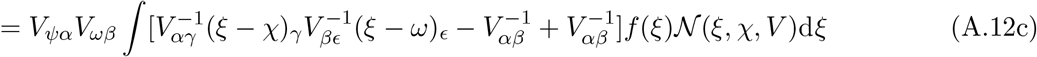

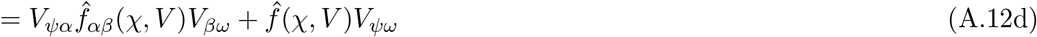

where we have used that *V* is symmetric so that *V_ψω_* = *V_ωψ_*.

Assuming that we have a normally distributed trait-density distribution 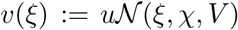 with total density *u*, mean *χ*, and variance-covariance *V* we can use the above results in turn to calculate three integrals that will appear in the derivation of the moment equations:

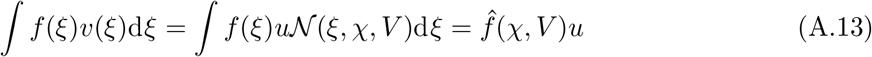

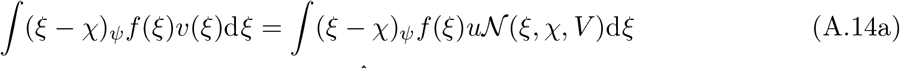

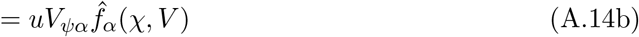

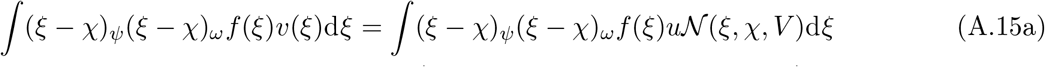

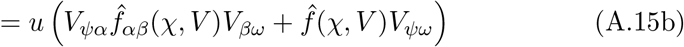

In the trait-space equations, we also need to evaluate the integrals over the normally distributed mutation kernels. In total, three such integrals will need to be evaluated, the first of which is:

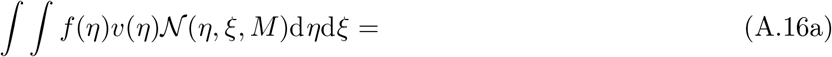

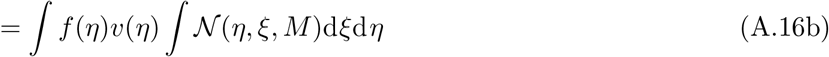

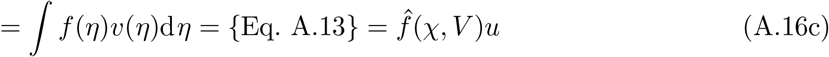

The second integral is

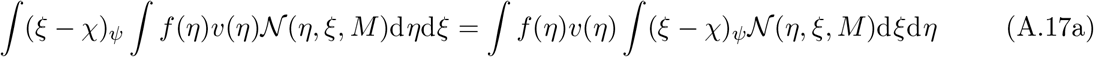

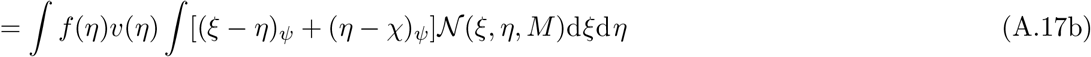

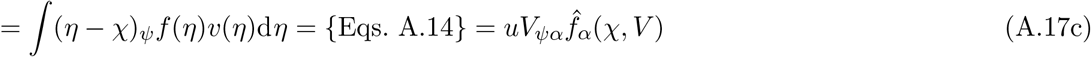

The third integral is

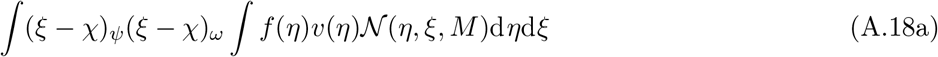

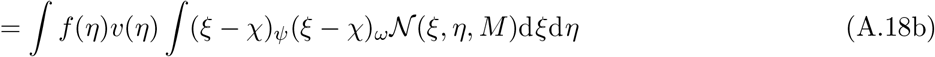

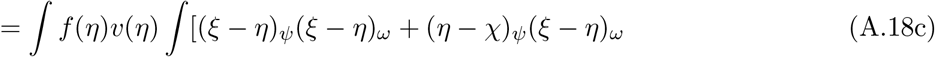

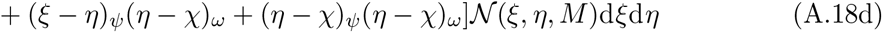

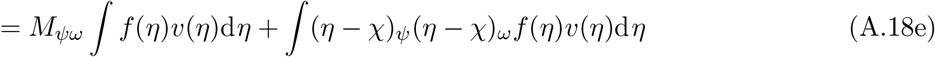

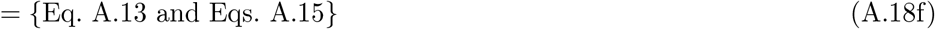

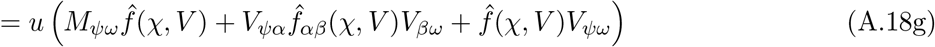

Recall that the trait-space equations are given by

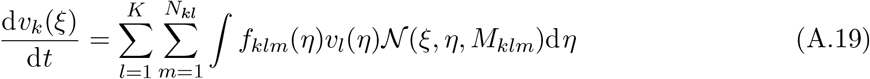

Here, the index *i* ∈ {1,…,*S*} pertains to species and *k,l* ∈ {1,…, *K*} pertain to classes. The index *m* ∈ {1,…,*N_kl_*} pertains to processes with focal class *k* and source class *l*. See main text for details. We now make the assumptions that the trait-density distribution in each class *v_k_*(*ξ*) can be decomposed into a sum of component distributions (species)

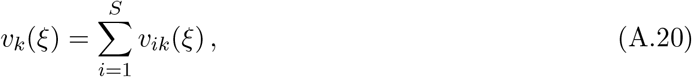

that each such component distribution can be approximated with a normal distribution

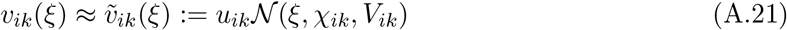

and that each species is reproductively isolated so that each species can be treated separately. Using the results derived above we can now calculate the moment equations for these trait-space equations.

<u>For *u*</u>:

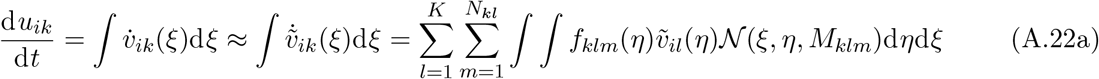

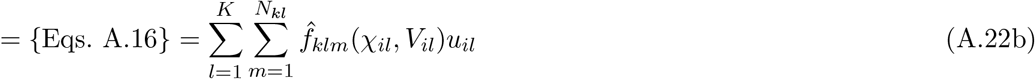

<u>For *χ*</u>:

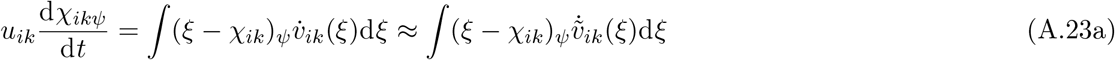

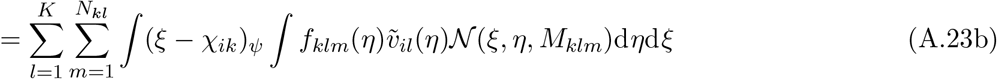

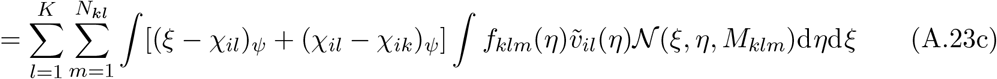

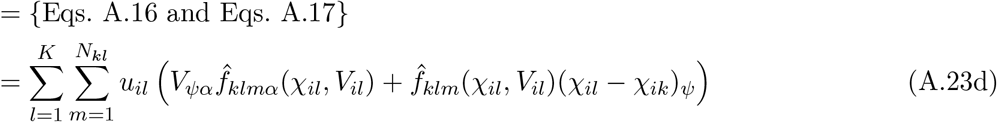

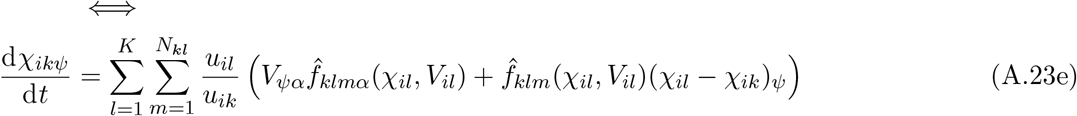

<u>For *V*</u>:

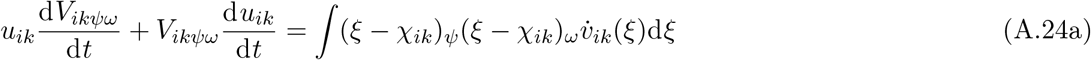

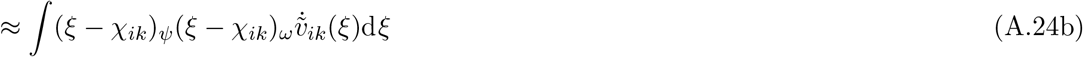

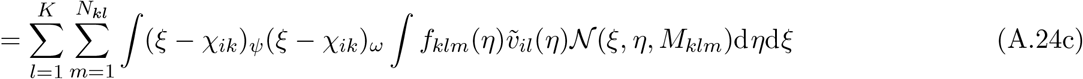

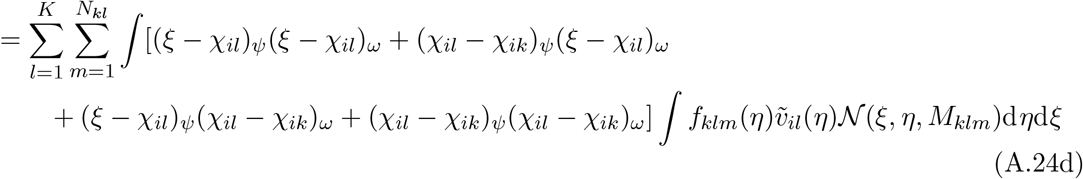

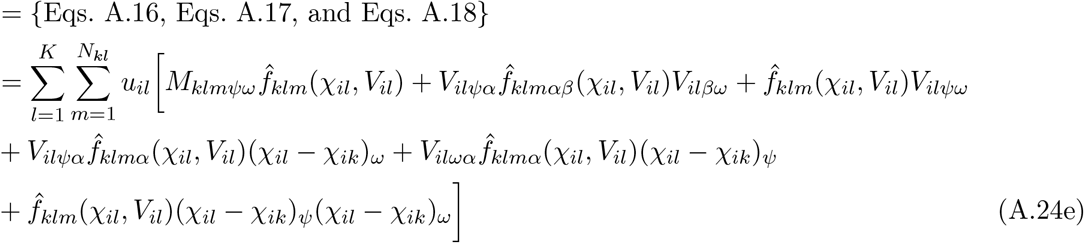

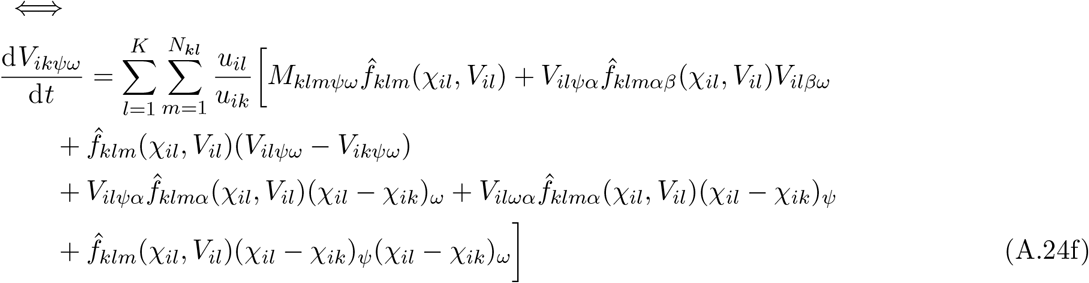

The complete moment equations for *S* species in *K* classes thus read

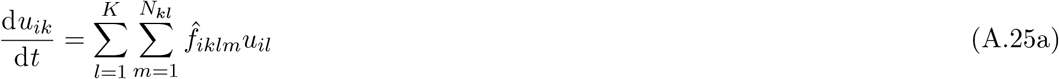

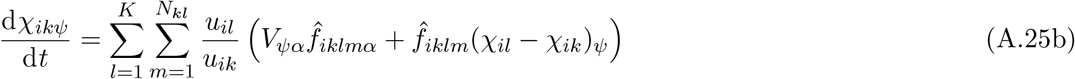

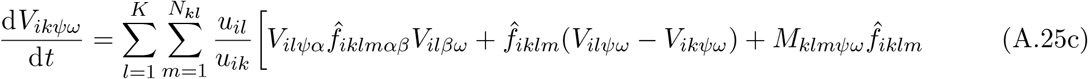

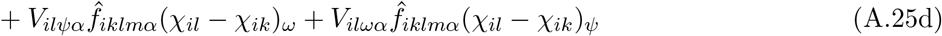

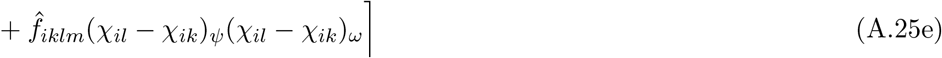

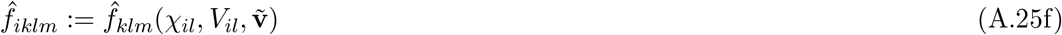

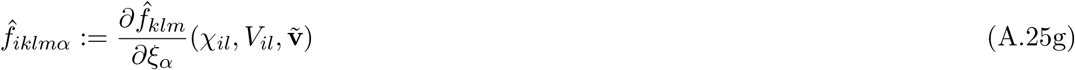

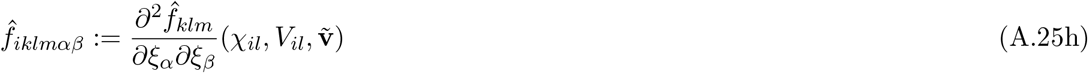

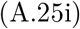

Here, we have omitted the explicit dependence of the process rates on its arguments for notational clarity. We have also added the dependence of the process rates 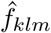 on the approximate traitdensity distributions 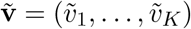 where 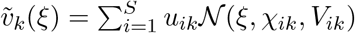. For convenience we did not include this dependence during the derivation, as their inclusion does not affect the derivation.

When the trait is scalar, i.e, *ξ* ∈ ℝ, these equations simplify to

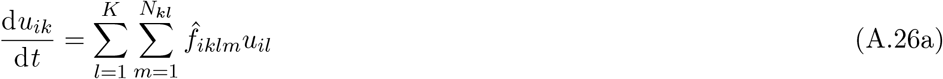

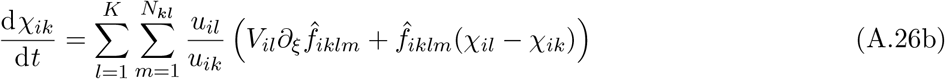

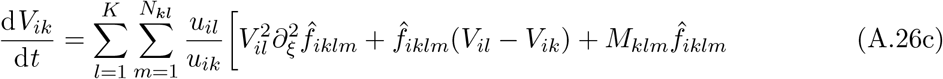

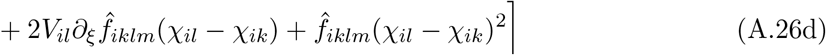

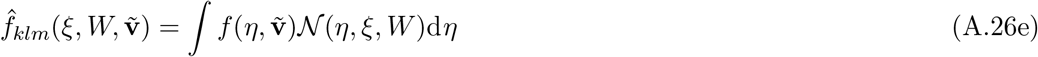

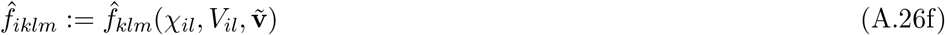

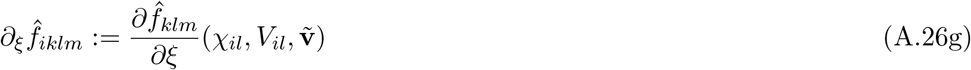

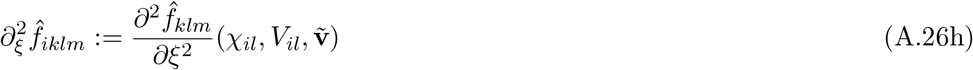

### Series solutions and Taylor approximations

The population per capita rates

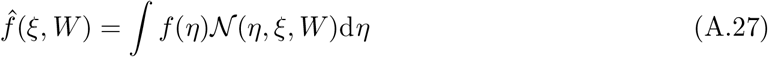

may not be analytically computable, and in such cases we may Taylor expand the process rates. To this end let *α^n^* denote an ordered set of indices such that

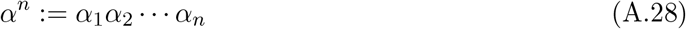

where each *a* indexes the trait-components. Thus, for example

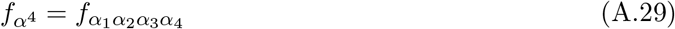

denotes the 4-tensor of mixed partial derivatives of order 4 of *f*. For a vector *ξ* in trait space we let

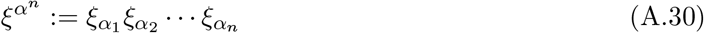

denote the outer product of *n* copies of the vector. For a matrix *W* acting on trait space we also let

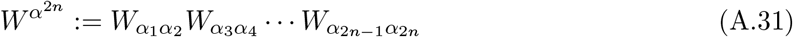

denote the outer product of *n* copies of the matrices.

We can now Taylor-expand

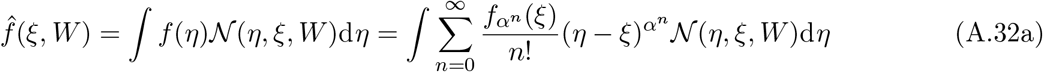

The central moments of a multivariate normal distribution are *n*-tensors and are given in e.g., Triantafyllopoulos (2002). However, when contracted against the fully symmetric tensor *f_α_n* these expressions simplify considerably so that

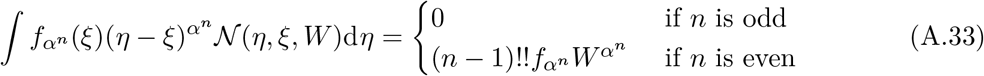

where “!!” is the double factorial, which is defined by

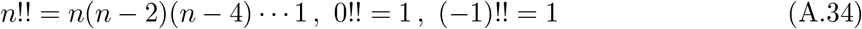

For the Taylor expansion we thus have

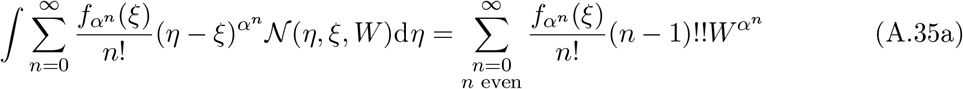

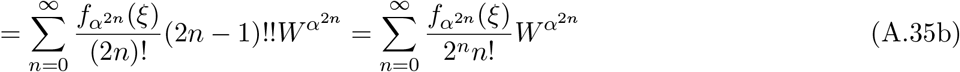

and hence

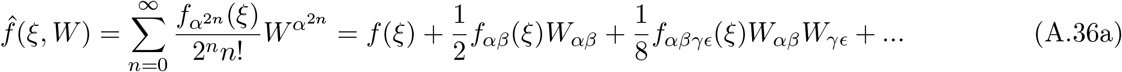

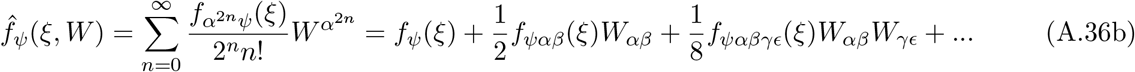

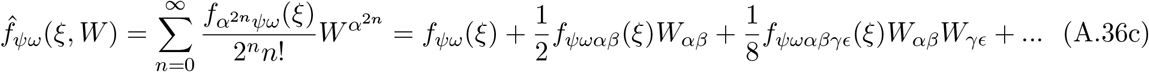

A Taylor expansion of order *n* then corresponds to truncating to derivatives of *f* with no higher order than *n*, so that e.g., a Taylor-approximation of order three would yield

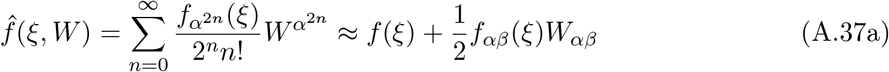

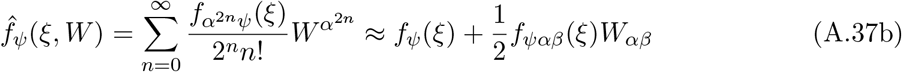

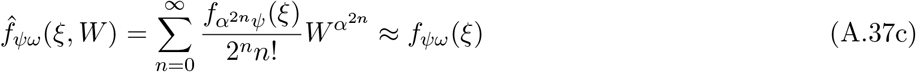

For a scalar trait, let *f*^(*n*)^ denote the *n*th derivative of *f* with respect to ξ. The expressions then simplify to

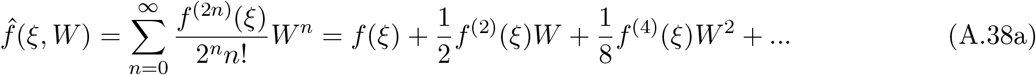

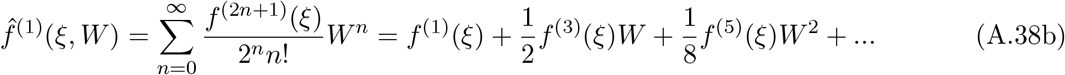

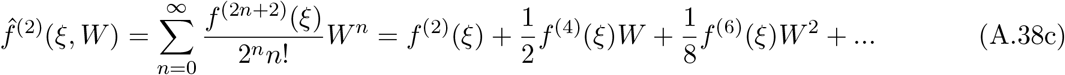

### Variable mutation variance-covariance matrices and mutation bias

For completeness we here presents result of more complicated mutation dynamics where the variance-covariance matrix is allowed to depend on the trait of the parent so that some phenotypes generate a wider array of offspring than others in trait space, and where mutations are not symmetric so that there is a bias of mutations in some direction in trait space.

First, we let the mutation variance-covariance matrix vary with trait so that *M*(*η*) is a positive-definite matrix for all *η*. The mutation kernel is then given by 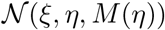. Note that for each fixed *η*, this is just the probability density function of a normal distribution over *ξ*.

Working through the problem in the same way as we did for constant mutation matrices (Eqs. A.16–A.17) we can then evaluate the three integrals used to account for mutation, the first of which is:

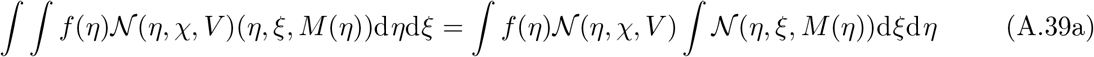

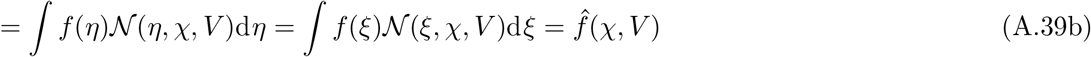

The second integral is

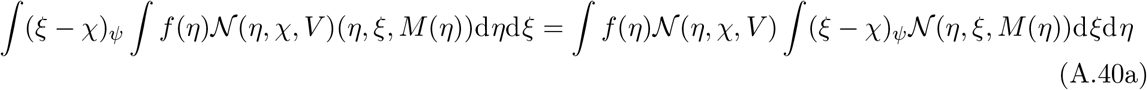

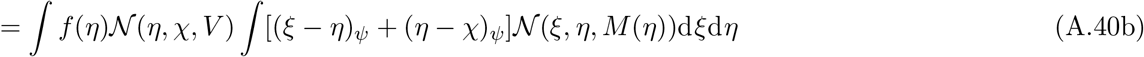

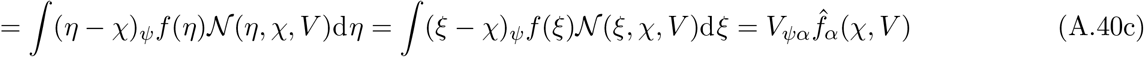

The third integral is

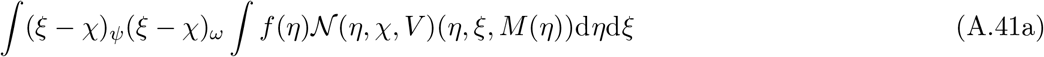

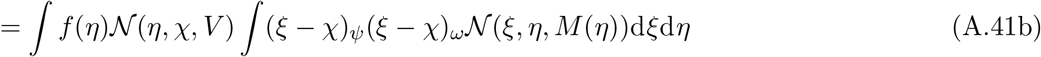

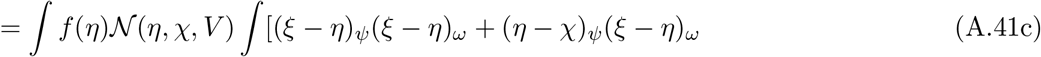

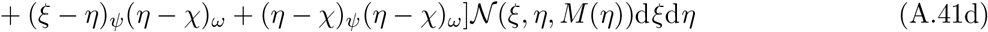

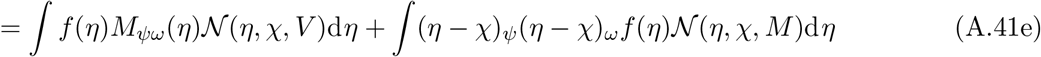

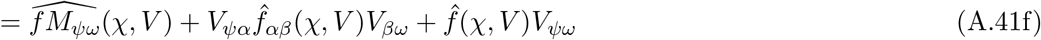

The mutation term in the moment equations when the mutation variance-covariance matrix is trait-dependent will thus read

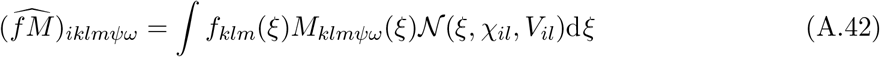

To second order we have

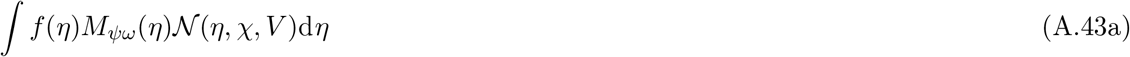

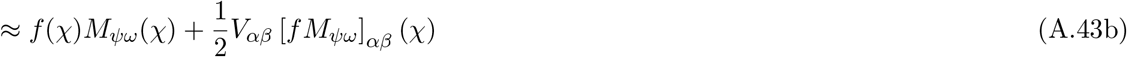

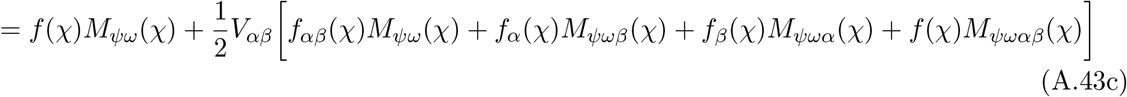

Apart from the term in the moment equations that add to variance-covariance through mutation, which is now given by Eq. A.42, the rest of the moment equations remain unchanged under the assumption that the variance-covariance matrix is trait dependent.

We now turn to the case where there is also mutation bias so that a parent with trait *η* on average produces offspring with trait *η* + *μ*(*η*) with variance-covariance *M*(*η*). Mathematically, this implies that a rate *f* produces new individuals with trait at a rate

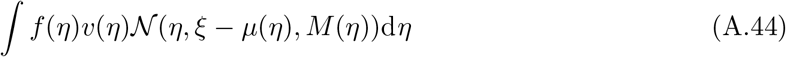

Once again, we can evaluate the mutation integrals, the first of which yields

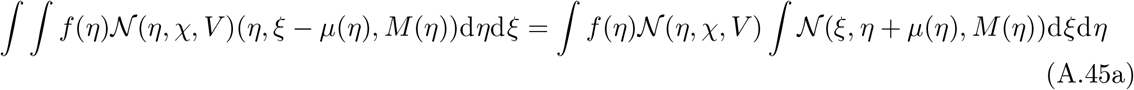

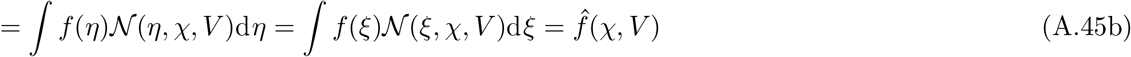

The second mutation integral is

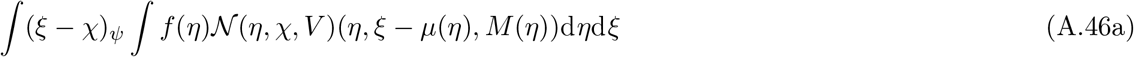

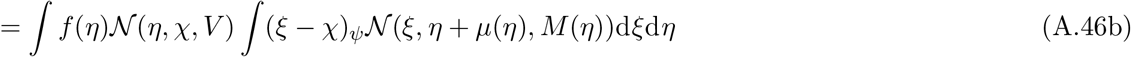

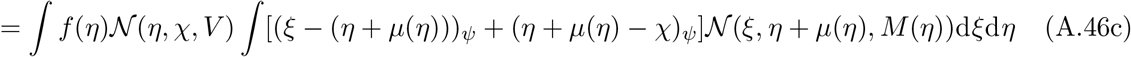

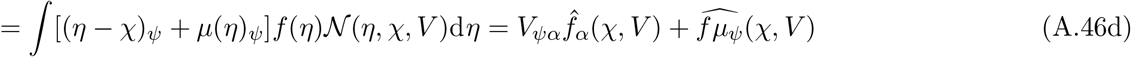

where

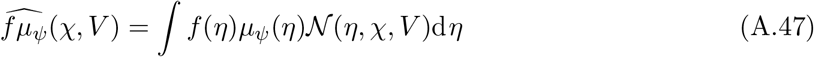

For the third mutation integral, let 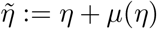. We then get

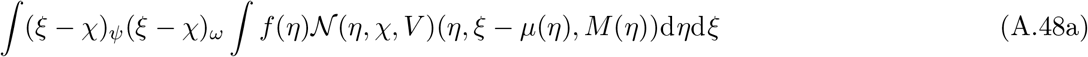

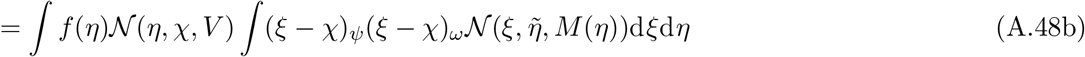

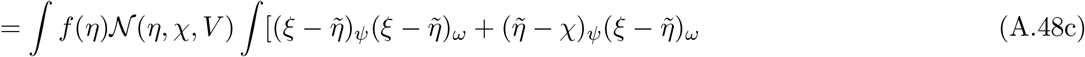

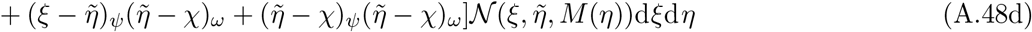

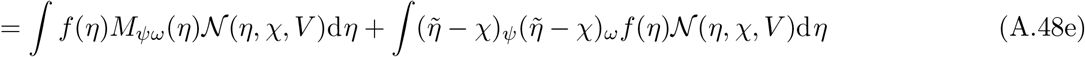

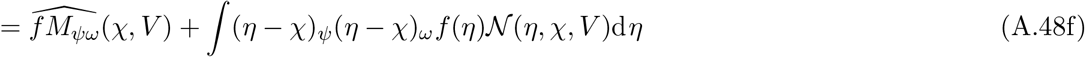

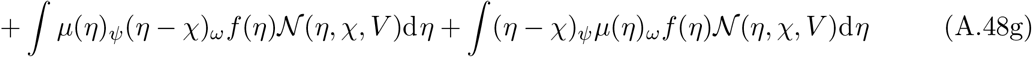

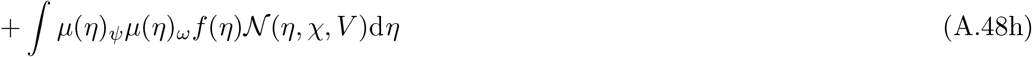

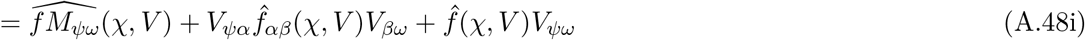

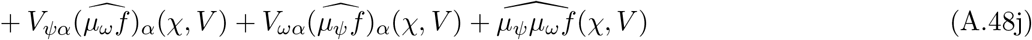

When inserted into the full moment equations where each process *f_klm_* has an attached mutation bias vector *μ_klm_*(*η*) for the class-structured community this yields the moment equations

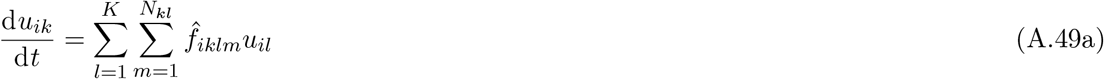

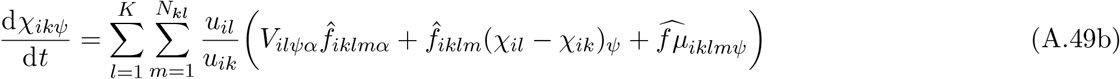

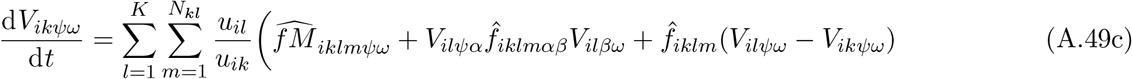

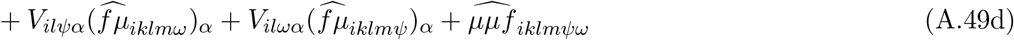

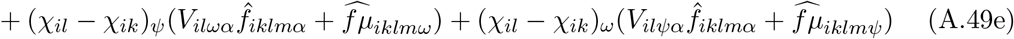

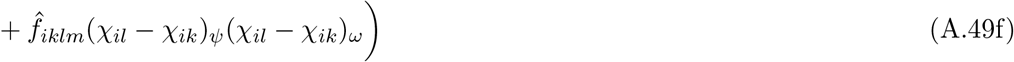

Here, all functions with subscript *iklm* are evaluated at *χ_il_* and *V_il_*.

In vector–matrix form these equations are written as

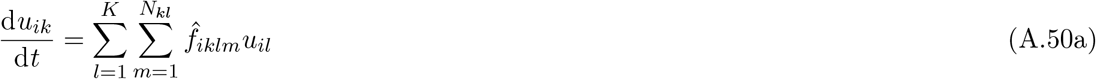

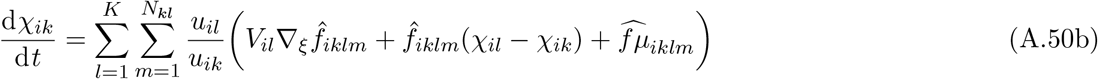

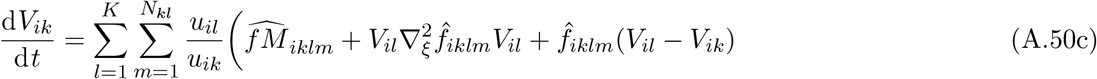

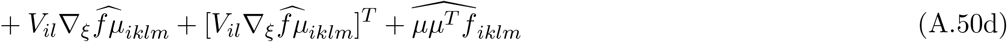

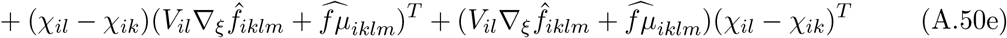

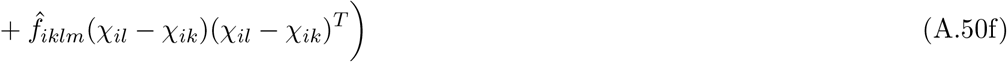

For scalar traits, these simplify somewhat to

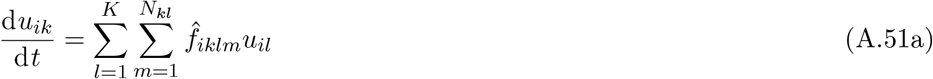

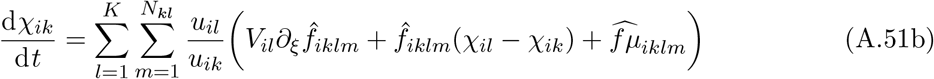

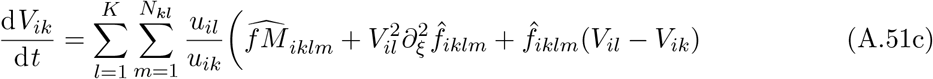

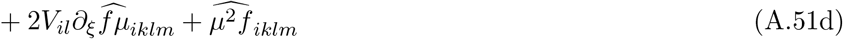

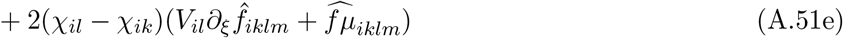

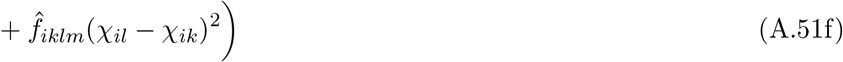

### Environmental variables

In our examples we have included non-evolving environmental variables. Such environmental variables include e.g., abiotic resources, toxins, or natural enemies whose traits can be considered fixed over the time-scale of interest. Here we present a brief account of how these are included in the traitspace and moment equations when the rate functions of the environmental variables depend linearly on the species’ trait-density distributions, as would be the case in classical resource-competition theory (Tilman, 1982) and modern niche theory (Chase and Leibold, 2003; Koffel et al., 2021).

Environmental variables may or may not follow the same class structure as the ecological communities. For example, in our two-patch model example, resources follow the same class structure as the species and are distributed among the patches, but in our stage-structured model the resources are unstructured. Furthermore, each environmental variable might have its own structure. Thus, let *E_jn_j__* be the density of environmental variable *j* in environmental class *n_j_*, with *N_E_* total environmental variables and *K_Ej_* environmental variable classes for environmental variable *j*. We let **E** denote the vector of all environmental variable densities in all environmental classes. We note here that it would be equally possible to simply enumerate all environmental variables *E*_1_, *E*_2_,… without loss of generality. However, for the times when the structure for the species mirrors that of the environmental variables (as in e.g., a patch model where each patch has, say, a resource and a natural enemy) structuring the environmental variables is useful.

Together with the trait-space equations for the ecological communities this yields the equation system

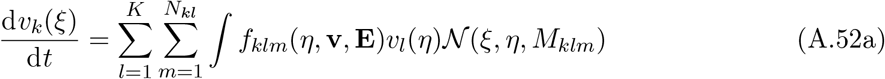

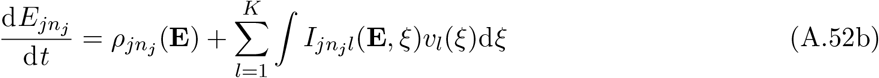

Here, *ρ_jn_j__* (**E**) represent the dynamics of environmental variable *j* in environmental class *n_j_* in the absence of any species, and *I_jn_j_l_*(**E**, *η*) is the impact of a an individual in class *l* with trait *η* on environmental variable *j* in environmental class *n_j_*.

As before, we now make the assumption that the trait-density distribution *v_l_*(*ξ*) can be approximated by a sum of normal distributions:

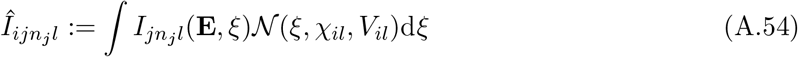

and, similarly to how we derived the moment equations, introduce the population-level per capita impact from a population with mean trait *χ_il_* and variance-covariance *V_il_*:

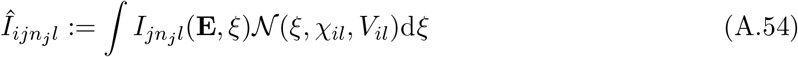

This can then be Taylor-expanded to arbitrary order. In particular, to second order this yields:

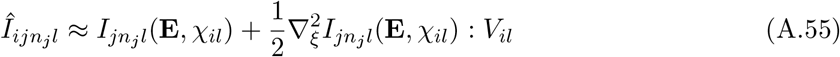

Thus, the moment equations including environmental variables read

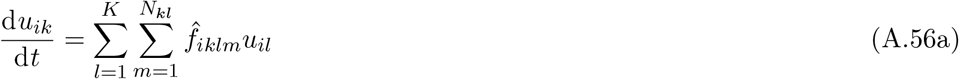

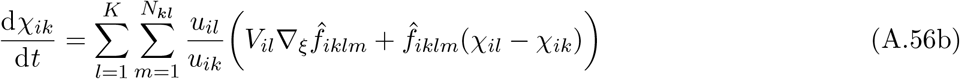

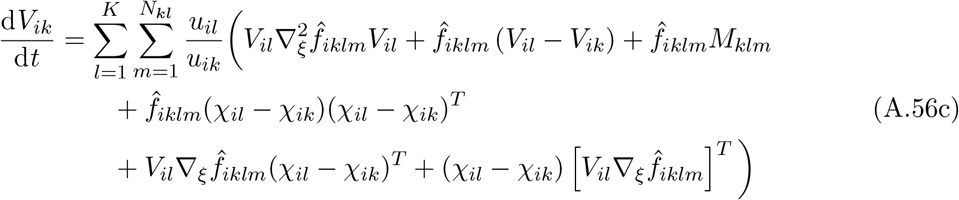

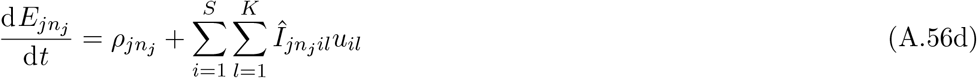

Here, the arguments of all rate functions have been suppressed for notational clarity, and functions are evaluated at *χ_il_, V_il_*, and **E**.

In Appendix C we use this general formulation to derive the rates of change for resources in a two-patch model and a stage-structured model.

### Explicit frequency-dependent dynamics

Though not part of any our example models, some models include explicit frequency-dependent dynamics. For example, consider our focal organism with trait-density distribution *v*(*ξ*) as the prey of a non-evolving predator with density *P* consuming the species with a type-II functional response with attack rate *a* and handling time *h*. If we now let the attack rates of the predators depend on the trait of the prey we have:

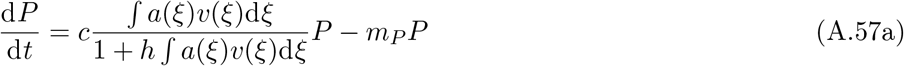

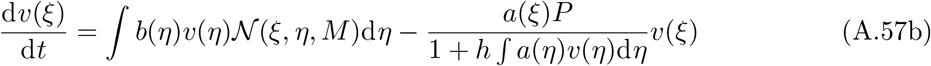

here, the mortality rate

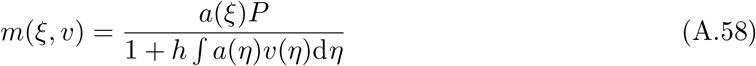

of the prey depends explicitly on the entire trait-density distribution of prey. However, under the assumption that the trait-density distribution *v*(*ξ*) can be approximated by a sum of normals, i.e.,

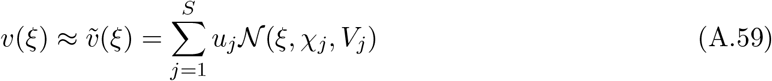

we may approximate the mortality rate by

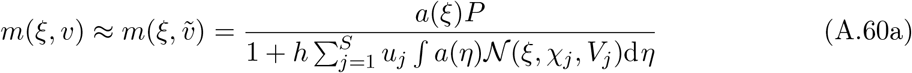

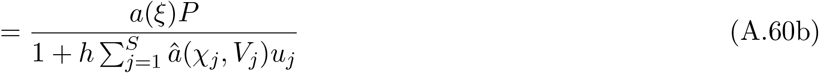

where

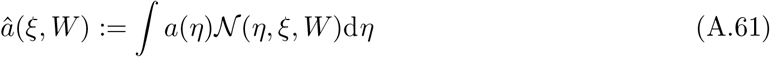

As for our general derivation, this integral may either be evaluated analytically if possible or approximated with a Taylor-expansion of *a*. Thus, as long as the explicit frequency dependence of any model comes in the form of functions of linear functionals of *v* this general recipe will always work. Typically, a linear functional *F* of *v* would take the form

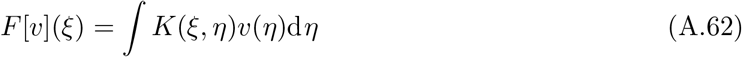

For example, in the predator–prey system above, *K*(*ξ,η*) = *α*(*η*).

Another example would be Lotka-Volterra dynamics (omitting mutations for brevity) where

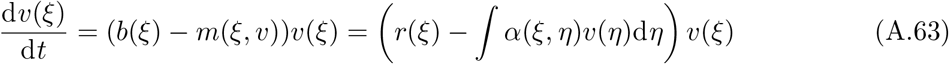

If we let

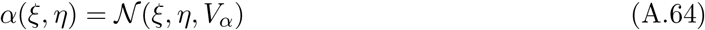

be the competition kernel, then, once again assuming that *v* is a sum of normal distributions we have

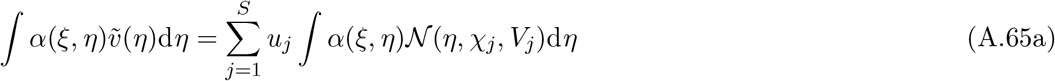

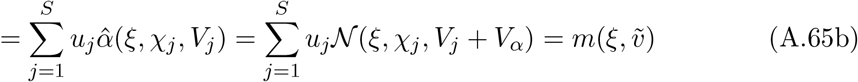

In this case we could solve the integral analytically, and furthermore, for our moment equations we can obtain

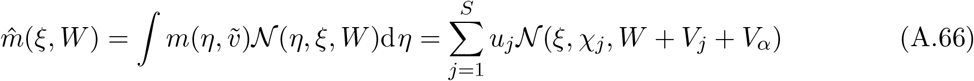

so that for species *i* we would have

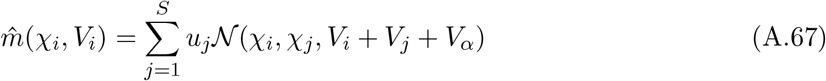

## Appendix B: Invasion heuristics

In this appendix we will derive ways of examining whether a community of species at eco-evolutionary equilibrium is closed to invasion by further species. We refer to these methods as heuristics since following these procedures cannot guarantee a complete closure to further invasions, and because the methodology is somewhat *ad hoc.* Nevertheless, in the examples we have tested (see the main text) these methods have so far proven to be highly accurate in the sense that they almost always produce communities with the same number of species for the moment equations as the trait-space equations produce peaks in eco-evolutionary equilibrium. The exceptions, as shown for the two-patch example in the main text primarily happens either when the number of species is ambiguous (a distribution with several peaks, but where the peaks are not well separated).

We shall follow the general ideas of adaptive dynamics (Metz et al. 1992; Dieckmann and Law 1996; Geritz et al. 1998; Dercole and Rinaldi 2008; Brännström et al. 2013) and work out two different ways of investigating when diversification into more species can take place. The first condition is local in trait space and known as an *evolutionary branching* where a species splits into two new species that subsequently diverge in trait space. The second condition treats global invasion analysis where the invader does not necessarily have a trait that is close to any resident species.

### Evolutionary branchings

We shall here work out a criterion for when a species characterized by the moment equations can be split into two new species locally in trait space in such a way that they subsequently diverge in their traits.

Consider a community of *S* species in *K* classes being in a stable eco-evolutionary equilibrium so that

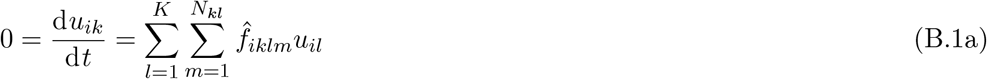

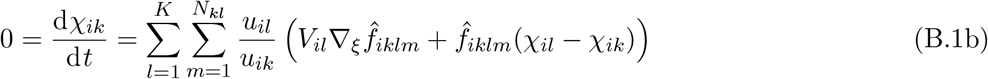

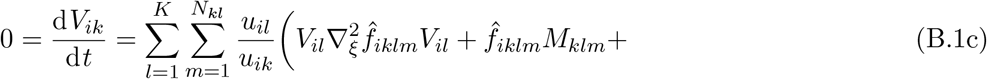

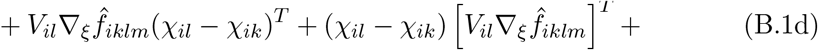

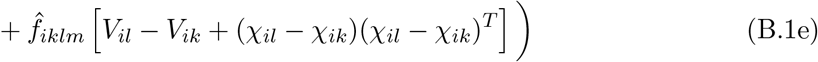

for all *i* ∈ {1,2,…,*S*} and *k* ∈ {1, 2,…, *K*}.

To test whether a given species *i* within this community can undergo an evolutionary branching we first split this species into two new species *i*_1_ and *i*_2_ with densities, mean-trait vectors, and variance-covariances given by

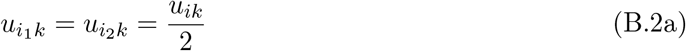

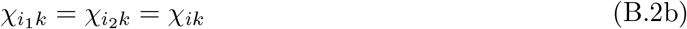

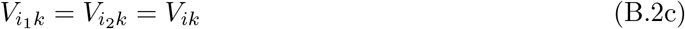

We then organize all the state variables, *u, χ*, and *V* into a vector *y,* so that the entries of *y* are the total densities, the mean trait components and the variance-covariance components for all species and classes in the split system. We let *F*(*y*) be the function given by the right-hand side of Eq. B.1 and linearize around the equilibrium *y*\*,

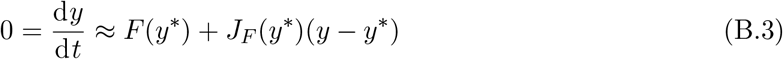

where *J_F_*(*y*\*) is the Jacobian matrix of *F* evaluated at *y*\*. Normally, in dynamical systems theory, we would say that the community with the split species is stable if the dominant eigenvalue (eigenvalue with largest real part) of the Jacobian matrix has a negative real part. However, here, due to the splitting, there will always be a neutral direction corresponding to shifting mass between species *i*_1_ and *i*_2_, so there will always exist a zero eigenvalue of the Jacobian. Thus, if the dominant eigenvalue is zero there is no branching. However, if there exists an eigenvalue with positive real part this indicates that the split-species equilibrium is unstable and that a branching in species *i* will take place.

Figure B.1 shows an evolutionary branching for the two-patch model (Box 1, main text). The blue species evolves to an eco-evolutionary equilibrium with substantial variance and significant local adaptation between the two patches. There, we split the species and determine the dominant eigenvalue *λ*_d_ as described above. For this example, the dominant eigenvalue is positive showing that the one-species equilibrium is unstable to the addition of a species. Let *ϕ* be the eigenvector associated with the dominant eigenvalue *λ*_d_. The components of the eigenvector are

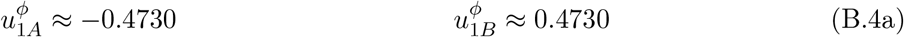

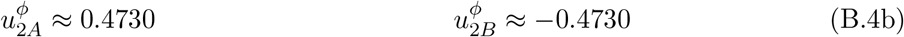

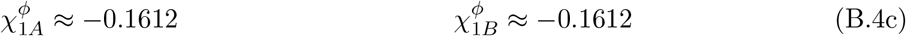

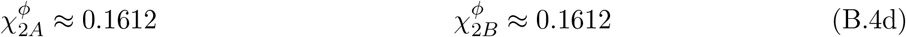

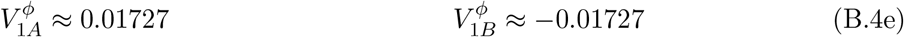

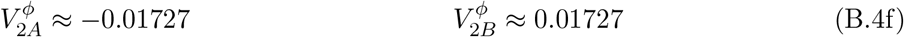

**Figure B.1:**
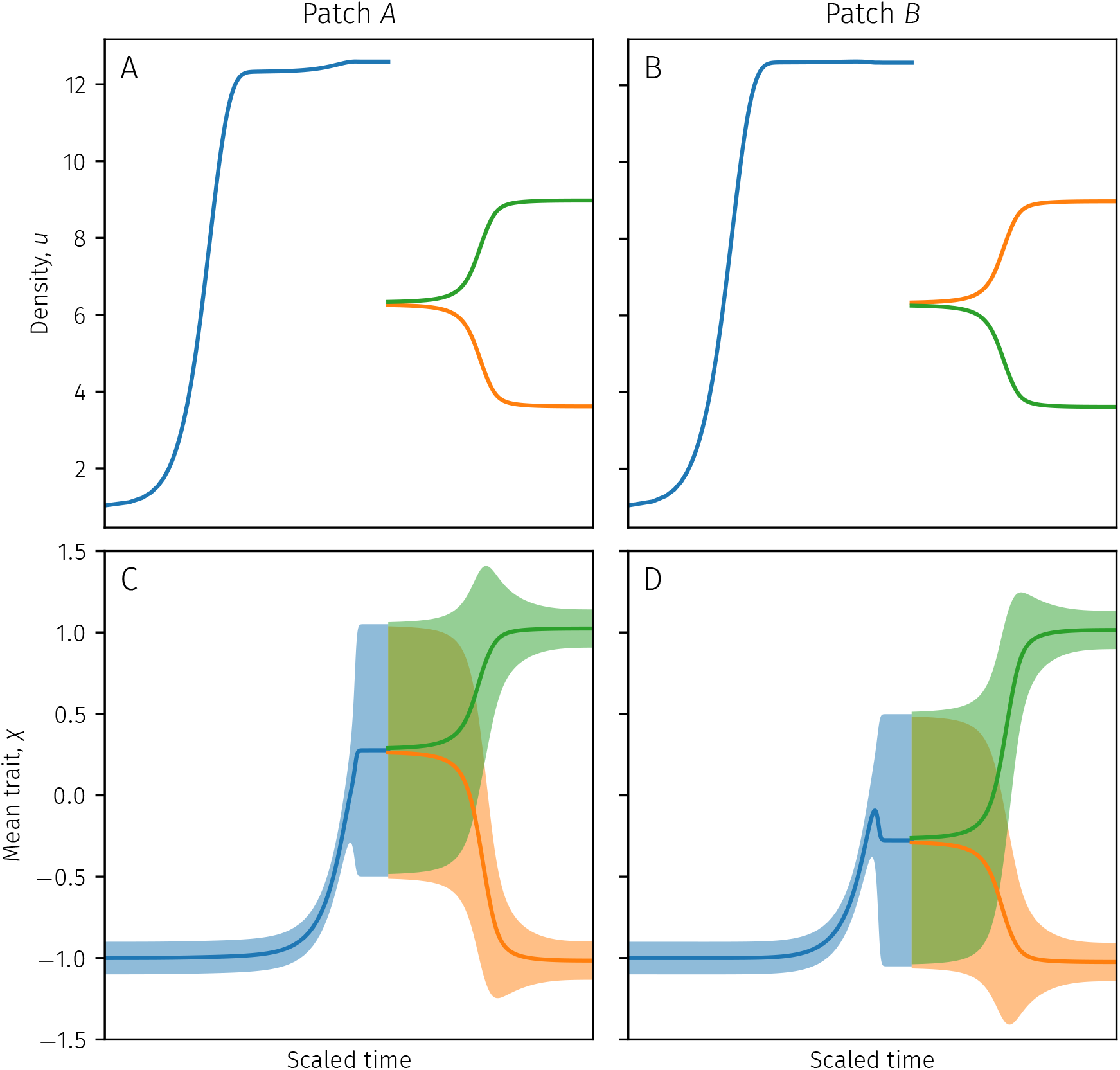
Illustration of branching procedure for the two-patch model (Box 1, main text). The panels depicts the densities (A–B) and mean traits (C–D) in the two-patch model on both patches where the left two panels depict patch *A* and the right two panels depict patch *B*. Panels (C–D) also depict one standard deviation around the mean as filled areas. The initial species (blue) evoloves to an eco-evolutionary equilibrium after which a branching takes place, the species is split, and the two new species (green and orange) once again evolve to an eco-evolutionary equilibrium. To illustrate the branching we have rescaled time non-uniformly to ensure the interesting aspects of the branching are clearly visible. **Parameter values**: *μ* = 0.1, *r* = 1, *K*_1*A*_ = 1, *K*_1*B*_ = 0.4, *K*_2*A*_ = 0.4, *K*_2*B*_ = 1, *c*_1_ = 1, *c*_2_ = 1, *α* = 0.5, *M* = 5 · 10^-6^, *d* = 8 · 10^-3^.

In other words, the unstable direction corresponds to one species becoming relatively more abundant on patch *A* and more resource one specialized, and one species becoming relatively more abundant on patch *B* and more resource two specialized. We construct the new two-species system after the branching by letting

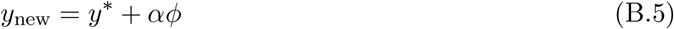

where *α* is some small number, for this example *α* = 0.05.

### Global invasion dynamics

Local branching in trait space will not always be sufficient to ensure that community is globally closed to further invasions. Here, we will work out a criterion for when an additional species with an arbitrary trait can be added to the community in such a way that this added species grows from being initially rare.

Let a discrete structured community of *S* species in *K* classes be in eco-evolutionary equilibrium so that

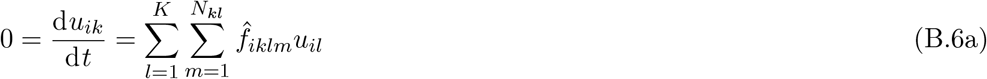

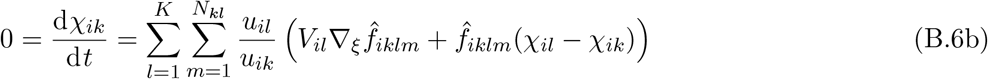

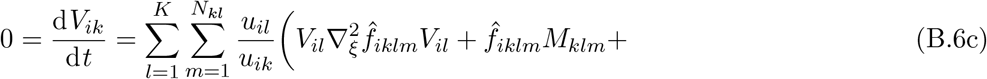

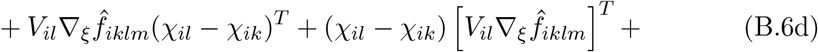

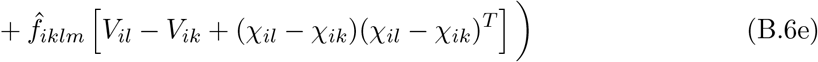

for all *i* ∈ {1, 2, …, *S*} and *k* ∈ {1, 2, …, *K*}.

We would now like to device a procedure to determine whether the resident community is closed to invasions. In analogy with adaptive dynamics (Metz et al. 1992; Dieckmann and Law 1996; Geritz et al. 1998; Dercole and Rinaldi 2008; Brännström et al. 2013) we will let a very rare invader enter this community, and try to determine whether an invader with a given trait can grow in the environment set by the resident. However, we also need to account for the internal dynamics of the invader, i.e., its mean traits and variance-covariances across classes. In principle, we would thus need to test for the ultimate invasion success of all invaders with all possible combinations of mean traits and variance-covariance matrices across all classes. This, however, presents two issues. First, the state space of all such combinations grows very quickly as the number of traits or number of classes increases making this approach infeasible. Second, since we often need to employ a Taylor approximation for the various rate functions governing the moment equations, we wish to avoid counting invaders with variances that are so large that their per capita growth rates can be spuriously positive due to a e.g., a positive second derivative. Thus, we take a two-stage approach for determining whether any successful invaders can be found. First, we assume that the variance-covariance of the invader is very small and let its mean traits evolve until equilibrium. Then, we let the variance-covariance matrices of the invader change until equilibrium as well, all while the invader is considered to be so rare as not to affect its environment. Finally, when the invader has reached an equilibrium in terms of mean traits and variance-covariances, we can calculate whether this invader will grows or decline in the environment set by the resident. Below, we will in turn make precise these notions and how to find any potential successful invaders. This invasion procedure was inspired by the invasion procedure used by Kremer and Klausmeier (2013).

For our first step, we assume that the invader is both very rare and that its variance-covariance is negligibly small. In this case, the term 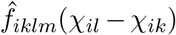 will dominate over the term 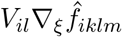 in Eq. B.6b, with the result being that the mean trait vector can be considered to be the same in each class (c.f., Wickman et al., 2017), making the situation analogous to adaptive dynamics. We can thus calculate the invasion fitness of the mutant with mean trait *χ*^inv^ as the dominant eigenvalue of the matrix *F*^0^, where the entries of *F*^0^ are given by

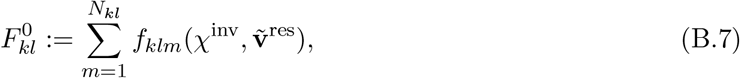

see e.g., Caswell (2001). This *adaptive-dynamics fitness landscape* describes how a mutant with negligible density and variance-covariance (and hence negligible local adaptation) would change over time in terms of mean trait, as the mutant would climb the fitness landscape to the peaks. For our second step, where we allow the variance-covariance matrices of the mutant to evolve, we thus need only consider such traits that correspond to the fitness peaks of this landscape (see Fig. 4 in the main text and Fig. B.2 for examples).

**Figure B.2:**
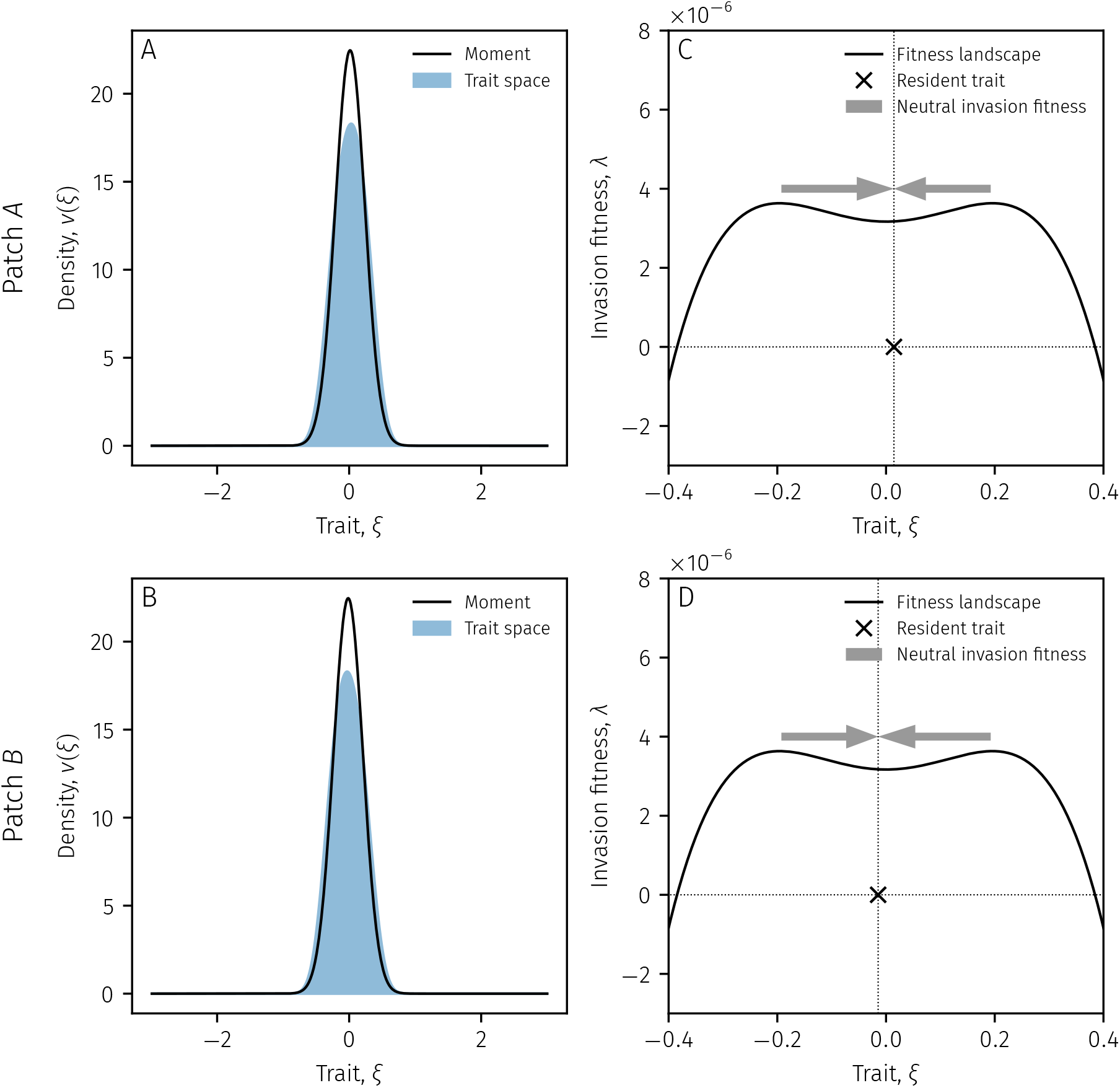
Illustration of a parameter set for the two-patch model (Box 1, main text) where the adaptive-dynamics fitness landscape does not predict qualitative features of the eco-evolutionarily stable community. **(A– B)** Tra it-dens ity d i str ibut i ons on patches *A* and *B* respectively i n eco-evoluti onary equil ibr i um of both the moment equati ons (black l nes) and tra t-space equat ons (blue areas). The moment equat ons capture the qual tat ve nature of the equ l br um, but is limited in its quantitative power due to the fact that the trait-space solution becomes platykurtic for this set of parameters. Comparing the moments on patch *A* we have 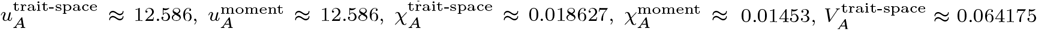, and 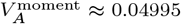. **(C–D)** Adaptive-dynamics fitness landscapes and invasion process on patches *A* and *B* respectively. The gray arrows illustrate the path of two invaders starting with near-zero variance at the fitness peaks and how their mean traits change over time as the variances and mean traits evolve over time for the invaders. The invaders end up with the same mean traits and variances as the resident and thus have neutral invasion fitness. The two-peaked fitness landscape implies that under zero-variance dynamics as in adaptive dynamics, the system would have a two-species eco-evolutionary equilibrium. Parameters are as in Fig. 3 in the main text except for *d* = 1.8 · 10^-2^.

There is however a problem that first needs to be overcome: The equations for the mean traits and variance-covariance matrices depend explicitly on the densities *u_ik_*. Crucially however, they only depend on the relative densities of the classes *u_il_/u_ik_* for some classes *k* and *l*. We will use this fact to formulate a set of invasion equations that allows the density distribution over the classes to vary without the total density of an invader across all classes changing over time.

First, we construct a matrix with entries 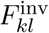 governing the per capita rates of an invader:

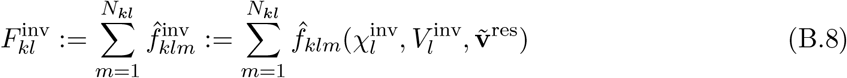

Given this notation, the density of a mutant would change over time according to

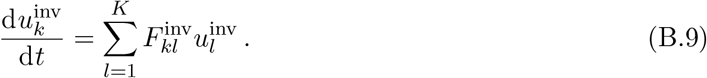

However, since we have assumed that the total density of the invader is very low, it will not affect *F*, and the densities of the invader will either grow towards infinity or decline towards zero, which will cause numerical problems. We can remedy this situation by tracking the relative densities across classes by changing *F* in such a way that it subtracts the total growth or decline across classes, while still allowing the invading mutant to assume the correct relative distribution among the classes. Thus, we let 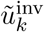 be the normalized frequency distribution among classes, i.e.,

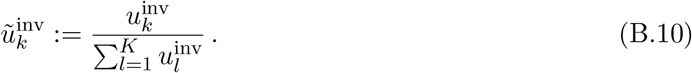

Furthermore, we let 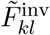 be the relative per capita growth matrix where the total growth is compensated for, i.e.,

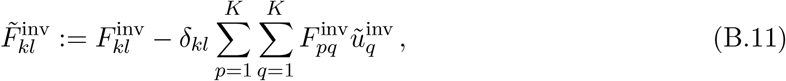

where *δ_kl_* is the Kronecker delta that equals one for *k* = *l* and zero otherwise. If we then let the densities of the invading mutant change according to

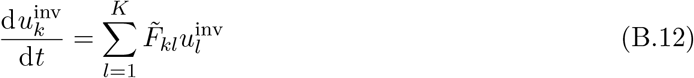

then the total density of 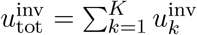 across classes, will not change over time, as can be seen by calculating

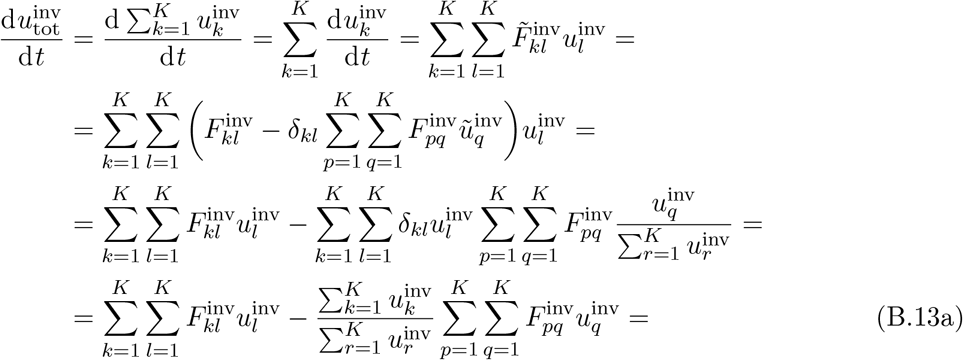

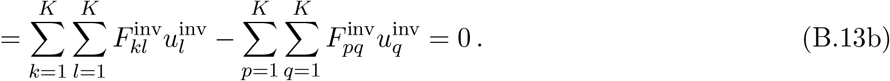

Having computed how to track the relative densities in each class, we may now carry out the second step of our invasion process, and evaluate the invasion fitness of the mutant at the end. The second step is computed as follows.

We pick an initial mean trait vector that maximizes the fitness landscape as described above, 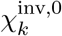, that is the same for each class *k*, and choose a variance-covariance matrix 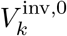 that is also the same for each class, without covariances and with some small variances for each trait. We let the initial density-distribution 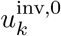 be the eigenvector associated with the eigenvalue of *F*^0^ at a fitness peak (the stable class distribution for zero variance-covariance), normalized so that 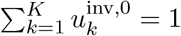. We then solve the equations

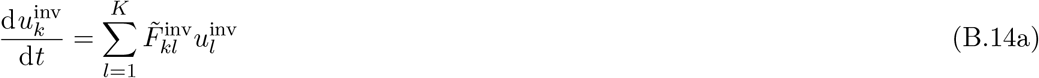

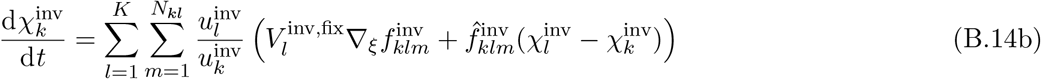

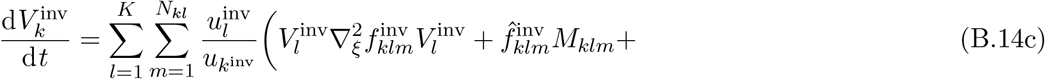

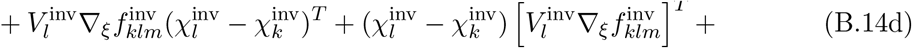

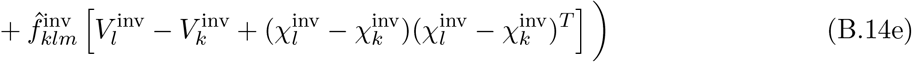

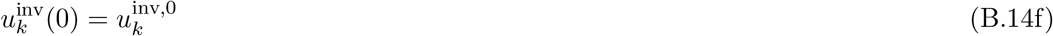

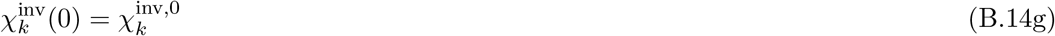

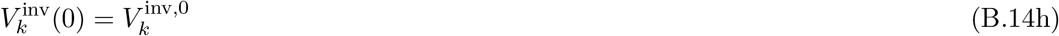

until they too are in equilibrium with 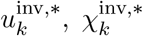, and 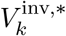 being the equilibrium densities, mean trait vectors, and variance-covariance matrices respectively.

Finally, we need to compute invasion fitness (i.e., the long-term exponential growth rate while rare) of the equilibrium invader. To this end, let 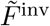 be the matrix that has 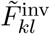 as its elements, and *u*^inv^ be the vector that has 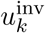 as its elements. Examining the equations for densities we get that

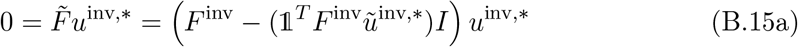

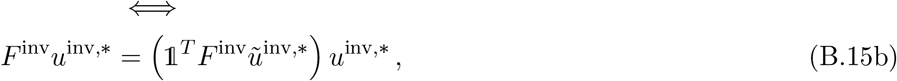

where 1 is a *K* × 1 vector having ones as all its elements and *I* is the *K × K* identity matrix.

From this we see that 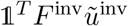 is an eigenvalue of the matrix *F*^inv^, and thus again in analogy with adaptive dynamics (see Caswell 2001 and Wickman et al. 2017 for structured populations), we can let

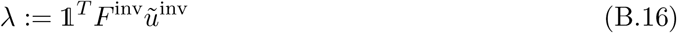

be the invasion fitness of the mutant.

We can now determine the ultimate fate of the mutant by looking at the sign of *λ*. If *λ* > 0, invasion fitness is positive, and the invader will successfully invade the resident community. If *λ* < 0, invasion fitness is negative and the invader will fail to invade the community. If *λ* = 0, the invasion fitness is neutral, and the invader will fail to invade the community. This last case typically occurs when the invading mutant ends up having the same mean traits and variance-covariances as one of the residents. This allows us to create invasion diagrams for structured populations, tracking the paths in trait space of invading mutants, and determining whether they will ultimately be able to successfully invade a community of residents. The process is illustrated in Fig. 4 in the main text.

We wish to emphasize that the zero-variance (adaptive-dynamics) fitness landscape is, in and of itself, insufficient to determine the invasion success of invaders. Figure B.2 illustrates a scenario where the fitness landscape would indicate a two-species equilibrium, but where the finite-variance case of the moment equations yields only one.

## Appendix C: Additional details of the example models

In this appendix we provide additional details of the example models presented in the main text, including details of how they were adapted from the generic equations, what numerical methods we used, and tables of parameter values used in numerical solutions.

### Two-patch resource-competition model

The trait-space equations described in the main text are given by

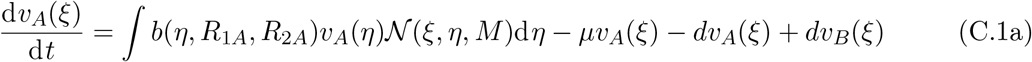

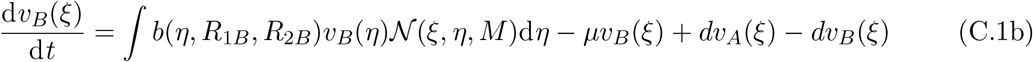

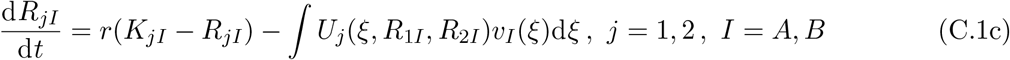

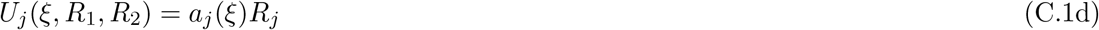

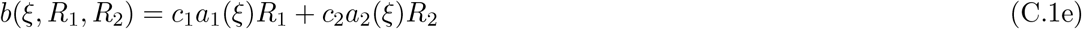

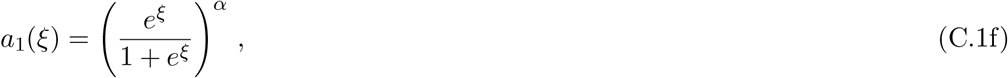

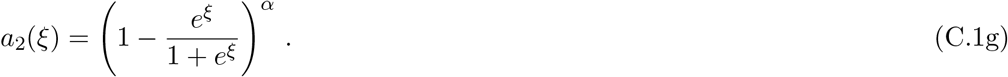

For our numerical solutions, especially for small mutation variances, the numerical solutions would require an inordinately high resolution in trait space, and so we approximate the mutation dynamics by a diffusion approximation (Kimura, 1965; Débarre et al., 2013)

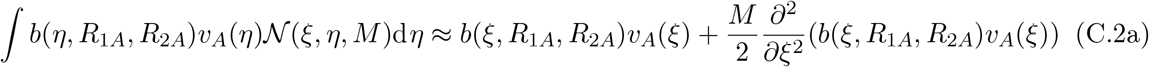

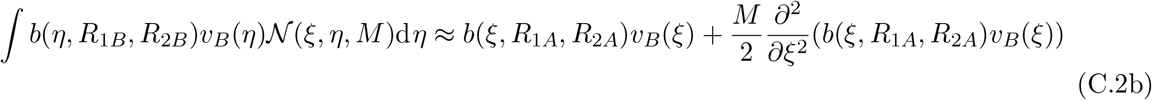

We then discretize trait space into 512 equidistant points between *ξ* = —5 and *ξ* = 5, and approximate the second derivative by a finite-difference approximation. On the edges, *ξ* = –5 and *ξ* = 5 we assume reflective boundary conditions, but this assumption is highly unlikely to affect results as the trait-density distributions are always very close to zero near the edges.

To get all eco-evolutionarily stable solutions of the trait-space equations depicted in Fig. 5 we run the discretized trait-space equations for 10^7^ time units for each combination of *d* and *M* using the `DifferentialEquations.jl` library (Rackauckas and Nie 2017) in Julia (Bezanson et al. 2017).

The moment equations were solved numerically using the same differential equation library. Comparisons between trait-space and moment equation solutions were made by evaluating the moment equation normal distribution sum on the same grid on which the trait-space equations were solved, and the integrals in trait space were then approximated as

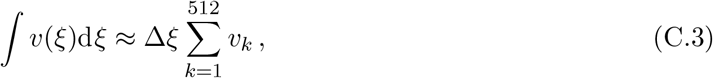

where Δ*ξ* was the spacing between grid points and *v_k_* the value of *v*(*ξ*) evaluated at the *k*:th grid point.

To get the moment equations for the resources *R_jI_* we use the generic equations for environmental variables derived in Appendix A and identify

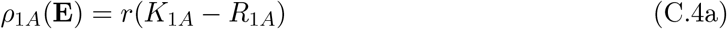

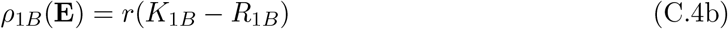

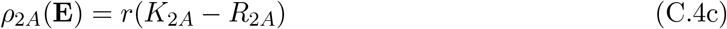

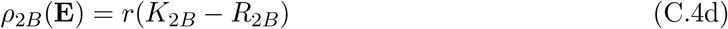

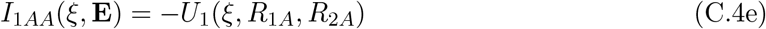

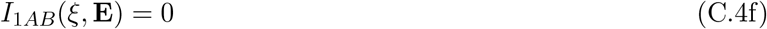

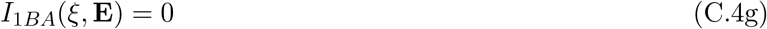

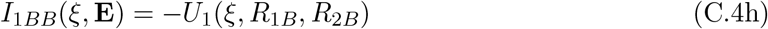

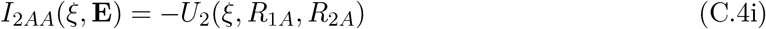

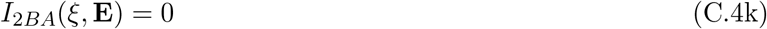

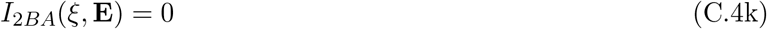

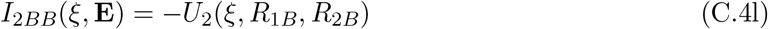

Comparison with the generic moment equations including environmental variables (Eq. A.56d) then yields that the resource dynamics in the moment equations will be

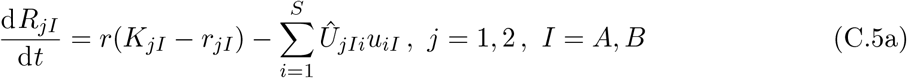

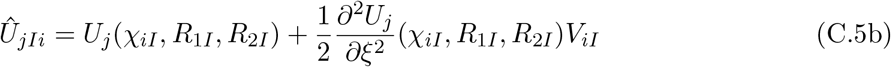

**Figure C.1:**
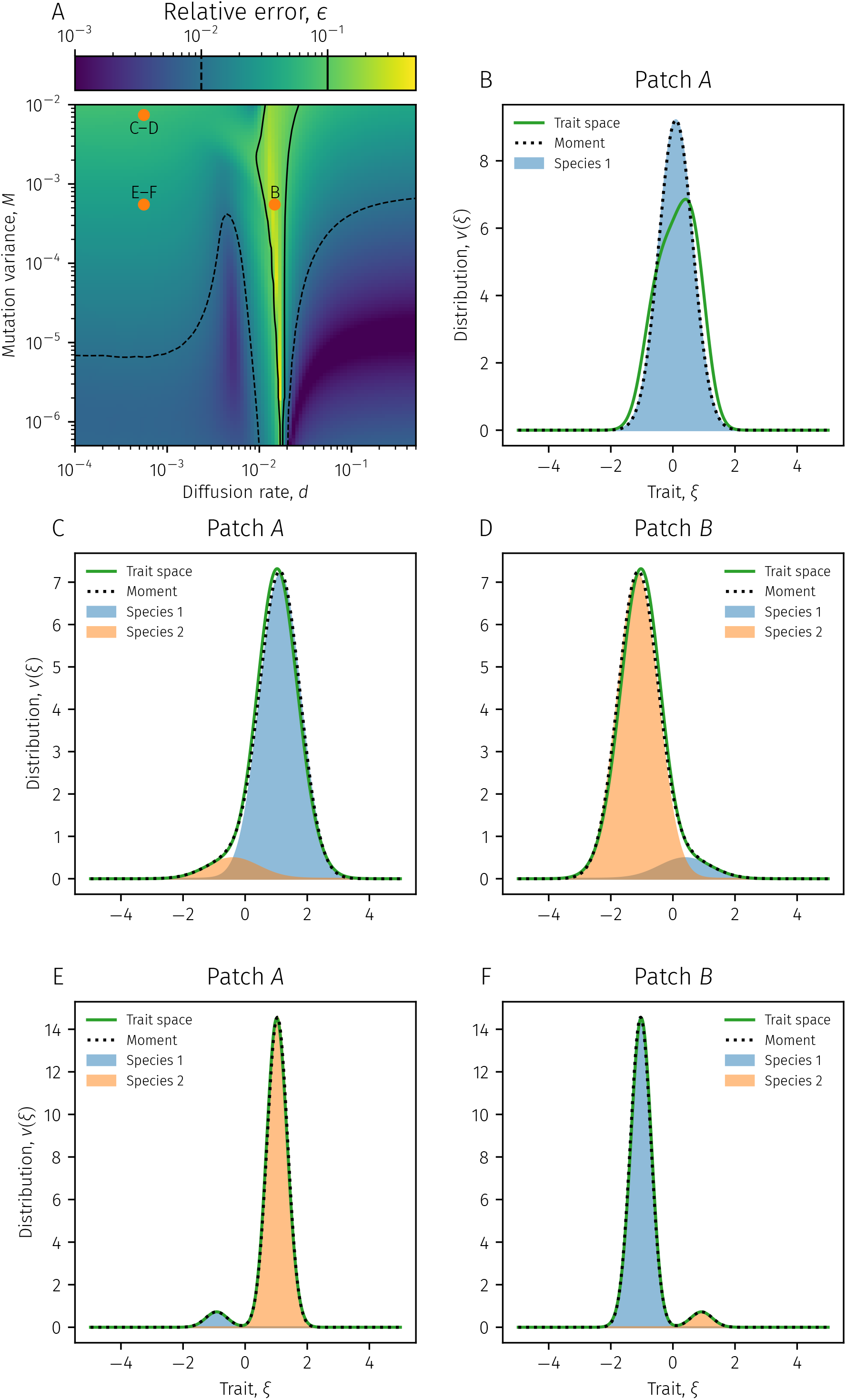
Examples of evolutionary cycling and trait-density distributions for high mutation rates in the two-patch model. **(A)** Reproduction of panel D of Fig. 5 in the main text showing the relative error between trait-space and moment solutions at eco-evolutionary equilibrium. The three dots show where in parameter space the examples shown in panels B, C–D and E–F are taken from. **(B)** Example of the moment-equation assembly procedure not accurately reprocuding the trait-density distribution of the trait-space equations. The equilibrium distribution of the trait-space equations (green) assumes a shape which is not easily represented as a sum of normal distribtuions and the moment equations (dotted black line), which here converge on a one-species equilibrium (blue area). Only patch A is depicted, as patch B has a near-identical, but reflected around *ξ* = 0, distribution. **(C–D)** Trait-space and moment solutions for high mutation variance and low dispersal on patches A and B respectively for panels B and C. Trait-space solutions are depicted as solid green lines, and moment solutions are depicted as dotted black lines. The two species that make up the moment solution are depicted as filled blue and orange areas. **(E–F)** Trait-space and moment solutions for intermediate mutation variance and low dispersal on patches A and B respectively for panels D and E. Trait-space solutions are depicted as solid green lines, and moment solutions are depicted as dotted black lines. The one species that make up the moment solution are depicted as filled blue areas.

**Figure C.2:**
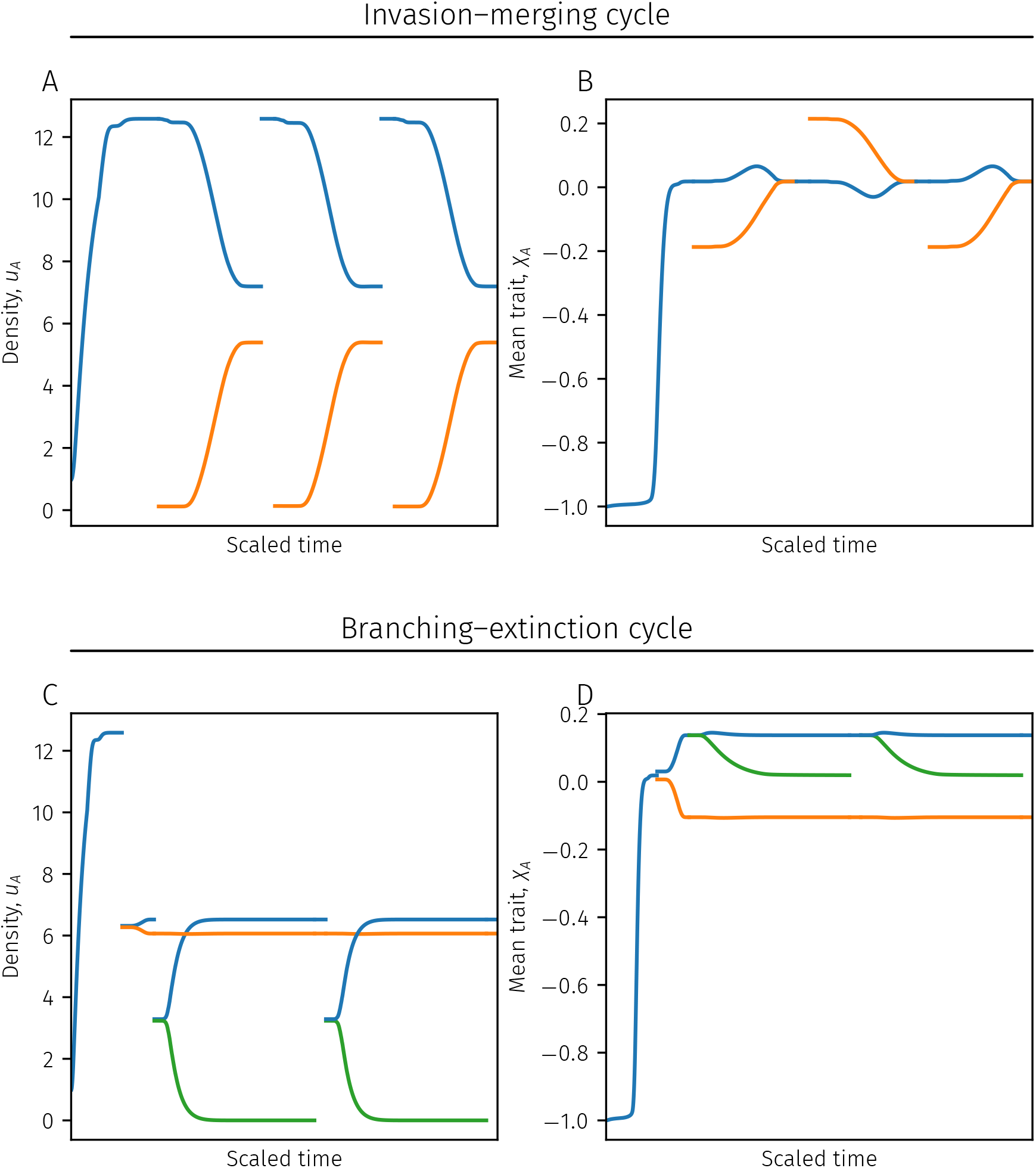
Examples of the two-patch model failing to converge to an eco-evolutionarily stable community. **(A– B)** Example of an invasion–merging cycle, where the densities (panel A) and mean traits (panel B) of the species on patch *A* are depicted. After settling onto a one-species equilibrium the resident species (blue) is invaded by a new species (orange). However, after successfully invading, the orange species evolves to the same mean trait as the blue species becoming equivalent. As the two species are equivalent, we merge them into one species and the cycle repeats. Paramter values: *d* = 0.017450183677616502, *M* = 3.3451854991893273 · 10^-6^. **(C–D)** Example of a branching–extinction cycle, where the densities (panel C) and mean traits (panel D) of the species on patch *A* are depicted. After settling onto a two-species equilibrium (blue and orange), one resident species (blue) undergoes a branching into two new species (blue and green). However, over time, one of the branches goes extinct (green) and is removed from the system, after which the cycle repeats. Paramter values: *d* = 0.017406558218422463, *M* = 3.3451854991893273 · 10^-6^. **Other parameter values:** *μ* = 0.1, *r* = 1, *K*_1*A*_ = 1, *K*_1*B*_ = 0.4, *K*_2*A*_ = 0.4, *K*_2*B*_ = 1, *c*_1_ = 1, *c*_2_ = 1, and *α* = 0.5.

### Stage-structured resource-competition model

The trait-space equations, adapted from De Roos et al. (2007), read

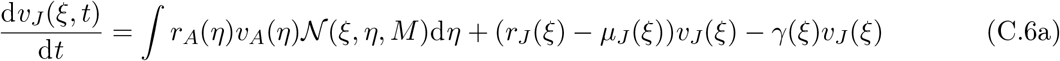

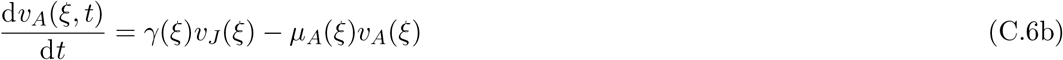

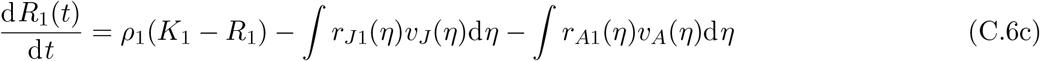

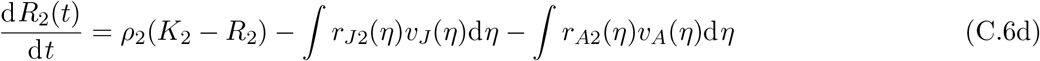

We can now compare these to the generic equations 2 in the main text, and identify the rate functions as

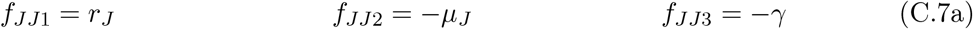

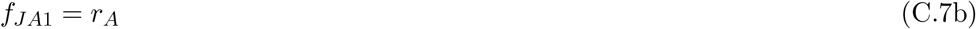

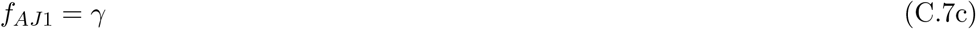

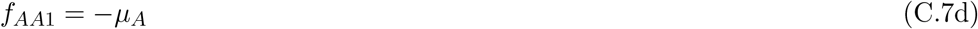

Mutations only occur in the transition from adults to juveniles, so *M_JA_* = *M* and the other mutation matrices are zero.

The necessary ingredients for the moment equations are now available, and we can insert the rate functions into Eqs. 5 in the main text to get the moment equations for the stage-structured model. Before we can write down the moment equations there we also need to identify what the renewal terms and resource impacts of the species will be. To that end, we compare with the trait-space equations including resources (Eqs. A.52) and identify

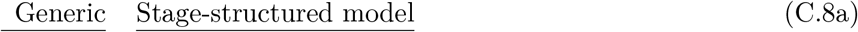

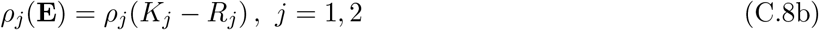

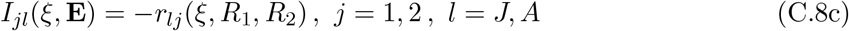

The moment equations for the resources will thus read

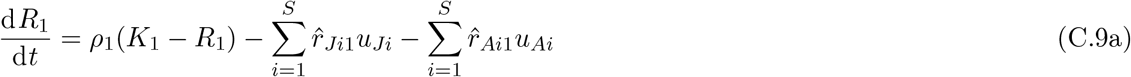

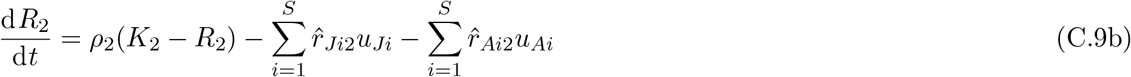

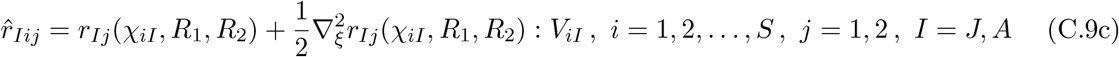

The resource uptake and stage-transition functions are given by

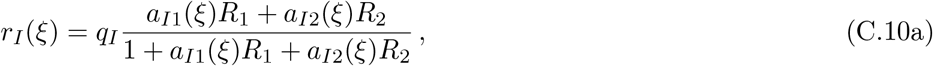

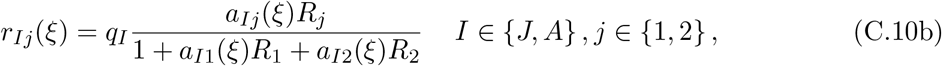

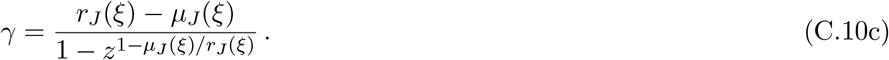

The maturation rate *γ* depends on the growth *r_J_*(*ξ*), the mortality *μ_J_* (*ξ*), and the size ratio between juveniles and adults *z*, see De Roos et al. (2007) for an explanation of how the specific form comes about. Here, *ξ* = (*ξ*_1_,*ξ*_2_) ∈ ℝ^2^, where *ξ*_1_ parameterizes a trade-off between the resource affinities and *ξ*_2_ parameterizes a trade-off between juvenile and adult mortality. Specifically,

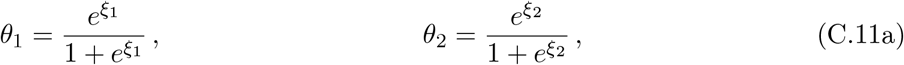

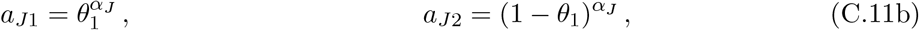

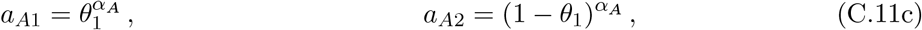

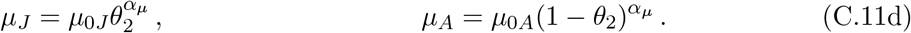

See Fig. C.3 for illustrations using the parameter values used in the numerical example. The parameters used in the numerical solutions are given in Table C.1.

**Figure C.3:**
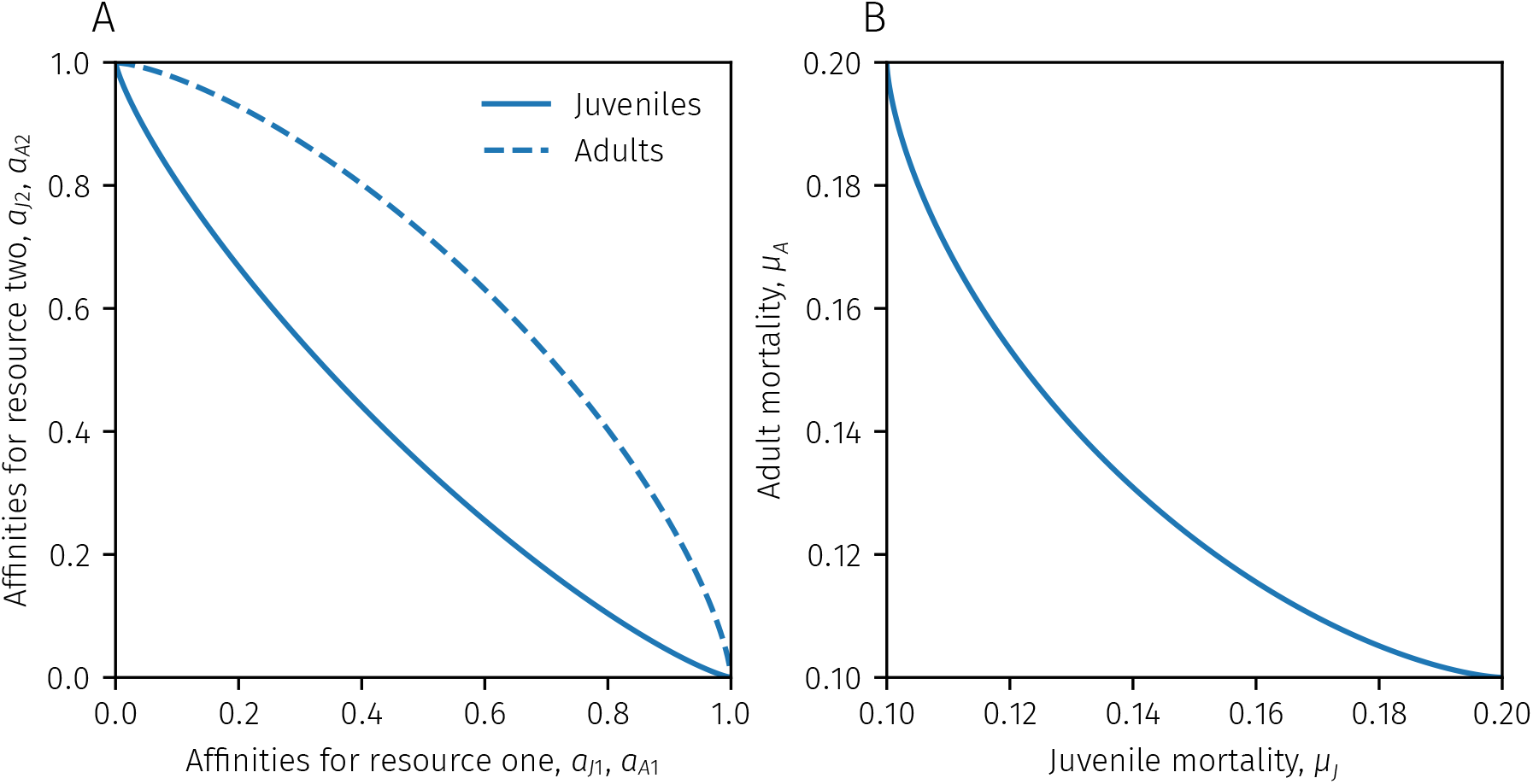
Trade-offs in the stage-structured model. **(A)** Trade-offs in resource affinities *a*_*J*1_ and *a*_*J*2_ for juveniles (solid line) and *a*_*A*1_ and *a*_*A*2_ for adults (dashed line). Juveniles experience a specialist-favoring trade-off, and adults experience a generalist-favoring trade-off. **(B)** Trade-off between juvenile and adult mortality. The trade-off between the mortalities is generalist-favoring. Equations for the trade-offs are given in Appendix C.

**Table C.1:** Parameter values used for numerical solutions of the stage-structured example.

| Symbol | Definition/Description | Value |
| --- | --- | --- |
| $\rho_1$ | Resource 1 renewal rate | 1.0 |
| $\rho_2$ | Resource 2 renewal rate | 1.0 |
| $K_1$ | Resource 1 supply | 1.0 |
| $K_2$ | Resource 2 supply | 1.0 |
| $q_J$ | Juvenile conversion efficiency | 1.0 |
| $q_A$ | Adult conversion efficiency | 1.0 |
| $z$ | Juvenile to adult size ratio | 0.3 |
| $\mu_{0J}$ | Juvenile base mortality | 0.1 |
| $\mu_{0A}$ | Adult base mortality | 0.1 |
| $\alpha_J$ | Juvenile resource trade-off shape parameter | 1.3 |
| $\alpha_A$ | Adult resource trade-off shape parameter | 0.7 |
| $\alpha_\mu$ | Juvenile–adult mortality trade-off shape parameter | 1.5 |
| $M$ | Mutation variance-covariance matrix | $10^{-5} \begin{bmatrix} 1.0 & 0.0 \\ 0.0 & 1.0 \end{bmatrix}$ |

Solving the trait-space equations for this system with these given parameters would require an unfeasible numerical resolution in trait space. However, since the mutation variance-covariance matrix is very small, without covariance, and with the same variances, we can approximate the birth/mutation integral with a diffusion term in the following way (Kimura, 1965; Débarre et al., 2013):

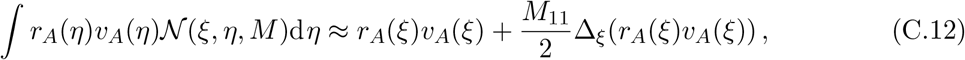

where 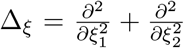 is the Laplacian in trait space. We discretize two-dimensional trait space into an equispaced grid of 256 × 256 points with *ξ*_1_, *ξ*_2_ ∈ [-4,4], and approximate the Laplacian with a central-difference approximation. We then used the `DifferentialEquations.jl` library (Rackauckas and Nie 2017) in Julia (Bezanson et al. 2017) to solve the semi-discretized ordinary differential equations to equilibrium. Both trait-space equations and moment equations were solved to equilibrium.

**Figure C.4:**
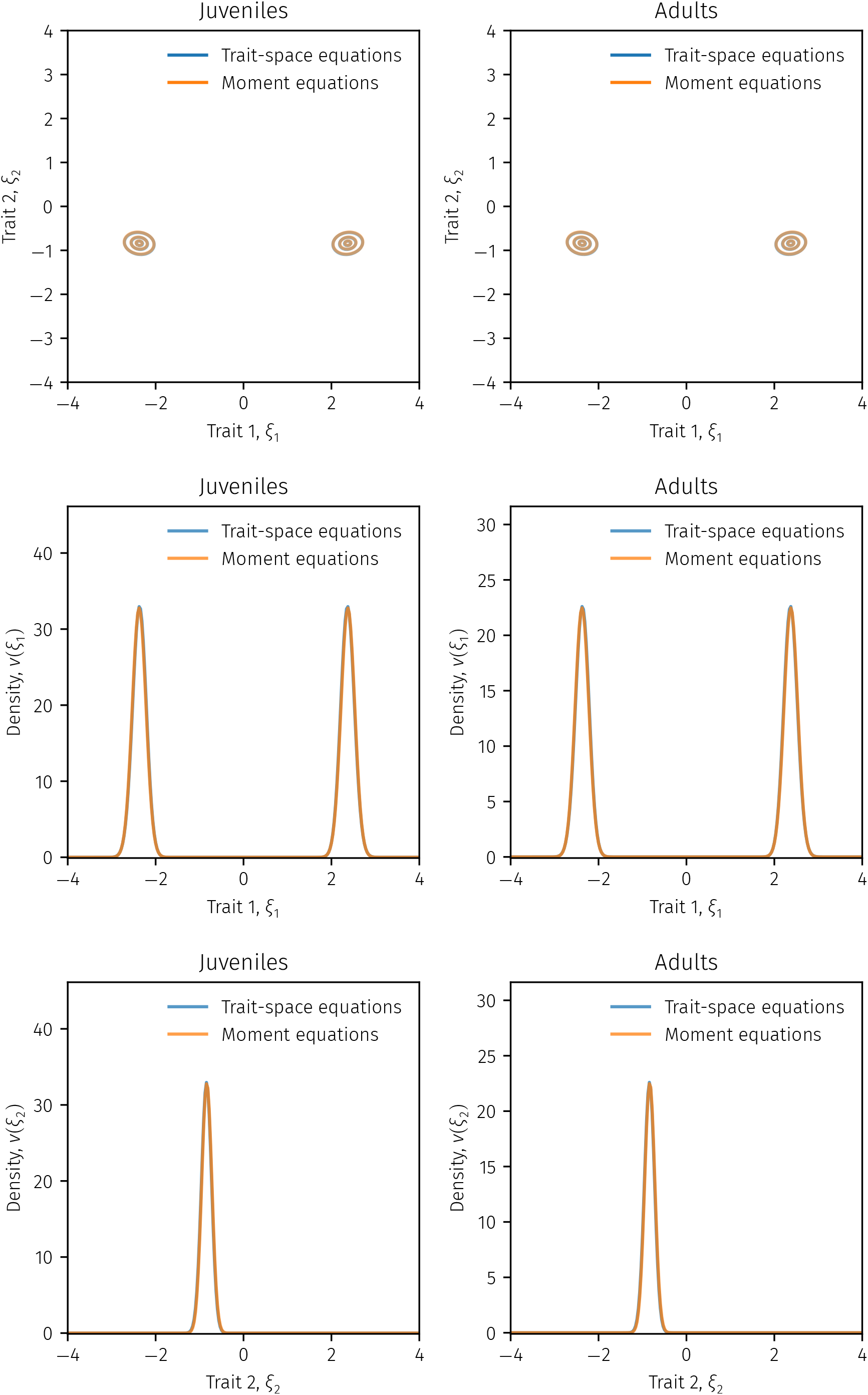
Comparison of trait-space equations and moment equations for the stage-structured model. Top row shows contour lines for the trait-space and moment equation solutions at eco-evolutionary equilibrium. The contour lines are very nearly on top of each other. The bottom four panels show slices in the i and 2 directions, where both slices are made across the global density maximum of the trait-space solution. The curves are very nearly on top of each other.

